# Bacterial SEAL domains undergo autoproteolysis and function in regulated intramembrane proteolysis

**DOI:** 10.1101/2023.06.27.546760

**Authors:** Anna P. Brogan, Cameron Habib, Samuel J. Hobbs, Philip J. Kranzusch, David Z. Rudner

**Affiliations:** Department of Microbiology Harvard Medical School Boston, MA 02115; Department of Cancer Immunology and Virology Dana-Farber Cancer Institute Boston, MA 02115

**Keywords:** Autoproteolysis, SEA domain, mechanotransduction, regulated intramembrane proteolysis (RIP), adhesion GPCR, cellulosomes

## Abstract

Gram-positive bacteria use SigI/RsgI-family sigma factor/anti-sigma factor pairs to sense and respond to cell wall defects and plant polysaccharides. In *Bacillus subtilis* this signal transduction pathway involves regulated intramembrane proteolysis (RIP) of the membrane-anchored anti-sigma factor RsgI. However, unlike most RIP signaling pathways, site-1 cleavage of RsgI on the extracytoplasmic side of the membrane is constitutive and the cleavage products remain stably associated, preventing intramembrane proteolysis. The regulated step in this pathway is their dissociation, which is hypothesized to involve mechanical force. Release of the ectodomain enables intramembrane cleavage by the RasP site-2 protease and activation of SigI. The constitutive site-1 protease has not been identified for any RsgI homolog. Here, we report that RsgI’s extracytoplasmic domain has structural and functional similarities to eukaryotic SEA domains that undergo autoproteolysis and have been implicated in mechanotransduction. We show that site-1 proteolysis in *B. subtilis* and Clostridial RsgI family members is mediated by enzyme-independent autoproteolysis of these SEA-like (SEAL) domains. Importantly, the site of proteolysis enables retention of the ectodomain through an undisrupted ß-sheet that spans the two cleavage products. Autoproteolysis can be abrogated by relief of conformational strain in the scissile loop, in a mechanism analogous to eukaryotic SEA domains. Collectively, our data support the model that RsgI-SigI signaling is mediated by mechanotransduction in a manner that has striking parallels with eukaryotic mechanotransducive signaling pathways.

**SIGNIFICANCE:** SEA domains are broadly conserved among eukaryotes but absent in bacteria. They are present on diverse membrane-anchored proteins some of which have been implicated in mechanotransducive signaling pathways. Many of these domains have been found to undergo autoproteolysis and remain noncovalently associated following cleavage. Their dissociation requires mechanical force. Here, we identify a family of bacterial SEA-like (SEAL) domains that arose independently from their eukaryotic counterparts but have structural and functional similarities. We show these SEAL domains autocleave and the cleavage products remain stably associated. Importantly, these domains are present on membrane-anchored anti-sigma factors that have been implicated in mechanotransduction pathways analogous to those in eukaryotes. Our findings suggest that bacterial and eukaryotic signaling systems have evolved a similar mechanism to transduce mechanical stimuli across the lipid bilayer.

## INTRODUCTION

Mechanotransduction is an emerging mode of signal transduction in which cells sense and respond to force stimuli. Well characterized examples include adhesion G Protein-Coupled Receptors (aGPCRs) and Notch receptors (1–3). In the case of aGPCRs, the extracellular domains of these receptors bind the extracellular matrix or surface ligands on a neighboring cell and shear force triggers aGPCR signaling. Similarly, Notch receptors bind membrane-anchored ligands on signal-sending cells. Endocytosis of the ligand-bound complex by the signal-sending cell generates a pulling force that activates Notch in the signal-receiving cell. In these and other examples, proteolysis plays a central role in mechanotransduction. aGPCRs undergo autoproteolysis in their extracellular domain but the cleaved fragments remain noncovalently associated. Mechanical force is hypothesized to pull the fragments apart revealing a membrane-tethered agonist that triggers GPCR signaling. In the case of Notch, the mechanical force exerted on its extracellular domain causes a conformational change that reveals a cleavage site for a membrane anchored protease. Ectodomain cleavage and release triggers intramembrane proteolysis by gamma-secretase and information transduction. Thus, this regulated intramembrane proteolysis (RIP) pathway is activated by mechanical force. In contrast to these well characterized eukaryotic systems, there are very few examples of bacterial mechanotransducive pathways and those that have been identified remain poorly understood. Our recent work on how the bacterium *Bacillus subtilis* responds to cell wall defects has uncovered a signaling pathway (4) that appears to combine features of aGPCR and Notch signaling.

Regulated intramembrane proteolysis (RIP) is a broadly conserved mechanism of information transduction that involves a two-step proteolytic pathway (5). In bacteria, the principal targets of RIP signaling are membrane-anchored anti-sigma factors that hold their cognate sigma factors inactive (6). External stimuli trigger “site-1” cleavage of the anti-sigma factor on the extracytoplasmic side of the membrane. A diverse set of bacterial proteases have been implicated in this regulated step. Ectodomain release allows the membrane-anchored portion of the anti-sigma factor access to the recessed interior of the S2P/RseP family of membrane-embedded “site-2” proteases (7, 8). Site-2 cleavage leads to release of the sigma factor and activation of gene expression. The *B. subtilis* RsgI anti-sigma factor is subject to RIP in response to cell wall defects (9) but unlike previously characterized bacterial RIP pathways, the activating step is not site- 1 cleavage. In this non-canonical RIP signaling pathway, site-1 cleavage appears to be constitutive but the two cleavage products remain associated. Activation of intramembrane proteolysis is mediated by their dissociation, which is hypothesized to involve a signal that generates mechanical force (4). Site-2 cleavage of RsgI releases the cognate sigma factor SigI that, in turn, activates genes involved in peptidoglycan (PG) biogenesis (10–12).

Here, we report that the extracytoplasmic domain of RsgI has structural and functional similarities to eukaryotic sea urchin sperm protein, enterokinase, agrin (SEA) domains (13). Intriguingly, many eukaryotic SEA domains undergo autoproteolysis and a subset have been implicated in mechanotransduction (14–17). We show that site-1 cleavage of *B. subtilis* RsgI and Clostridial homologs is mediated by enzyme-independent autoproteolysis in these SEA-like (SEAL) domains. Importantly, the site of proteolysis enables retention of the ectodomain through an undisrupted ß-sheet that spans the two cleavage products, similar to what has been observed for aGPCRs (18). Furthermore, we show that autoproteolysis can be abrogated by relief of conformational strain in the scissile loop, in a mechanism analogous to eukaryotic SEA domains (19, 20). Intriguingly, SEAL domains are present throughout Firmicutes where they are fused to anti-sigma factors as well as diverse extracytoplasmic domains. We demonstrate that several of these SEAL domains also undergo autoproteolysis at a conserved site and the cleaved fragments remain stably associated. Collectively, our data define a family of bacterial SEAL domains that arose independently from their eukaryotic counterparts and argue that RsgI-SigI signaling is mediated by mechanotransduction in a manner that is similar to eukaryotic mechanotransducive signaling systems.

## RESULTS

### The AlphaFold model of RsgI’s juxtamembrane domain is structurally similar to SEA domains

Previous genetic screens in *B. subtilis* that implicated the S2P/RseP family member RasP in the SigI-RsgI signaling pathway failed to identify a site-1 protease (4, 9). Attempts to identify this protease using targeted disruption of known secreted and membrane-anchored extracytoplasmic proteases were unsuccessful. Instead, our analysis of the AlphaFold2-predicted structure (21) of RsgI led us to discover that constitutive site-1 proteolysis is mediated by autoproteolysis. The predicted structure of RsgI contains an N-terminal cytoplasmic anti-sigma factor domain, a transmembrane segment, and an extracytoplasmic juxtamembrane domain (JMD) followed by a long intrinsically disordered region (IDR) (**Fig. 1A**). The RsgI model has an overall average pLDDT of 72.97, but a very high average pLDDT (94.16) within the folded JMD (**Fig. S1**). We have previously shown that site-1 cleavage occurs within the JMD and that the IDR is important for signaling and could be responsible for the mechanical force that dissociates the site-1 cleavage products (4). To investigate potential mechanisms of site-1 proteolysis and ectodomain retention, we used the high-confidence model to search for structural homologs of the JMD in the Protein Data Bank.

**Figure 1.**
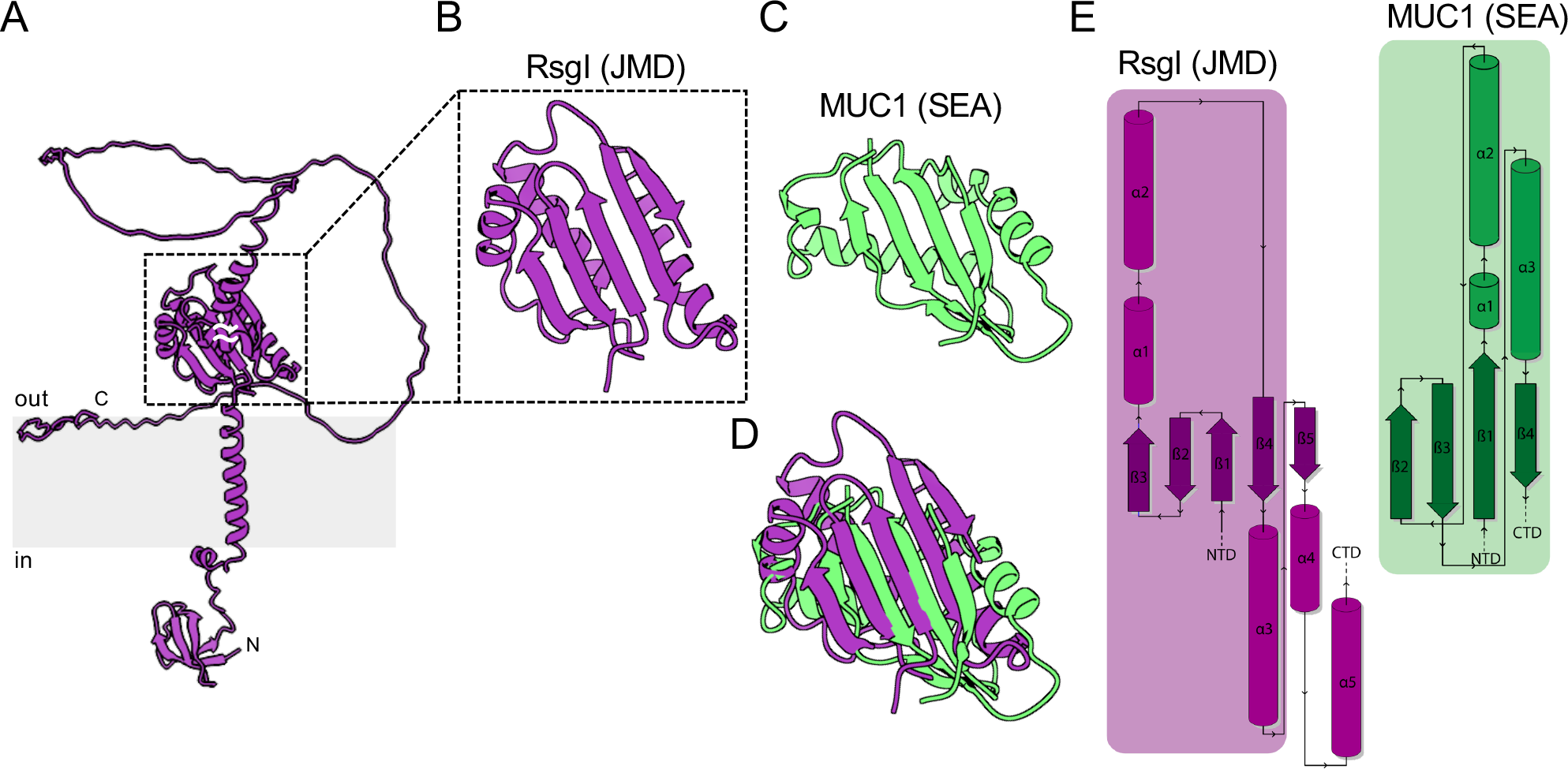
The Alphafold2-predicted structure of RsgI’s juxtamembrane domain has structural similarity to eukaryotic SEA domains. (A) AlphaFold2-predicted model of *B. subtilis* RsgI. The membrane bilayer is shown in grey. (B) Zoom in of the juxtamembrane domain (JMD). **(C)** Crystal structure of the SEA domain from *H. sapiens* Mucin-1 (MUC1) (pdb: 6bsb). **(D)** Structural alignment by TM-align of the AlphaFold2-predicted structure of RsgI’s JMD and the crystal structure of MUC1’s SEA domain. **(E)** Protein topology maps of RsgI’s JMD and the SEA domain in MUC1 as generated by PDBsum.

The top hit using the DALI server (22) was an N-terminal asparagine amidohydrolase that had modest structural similarity to RsgI’s JMD. Several other top hits were also enzymes, but none of their active sites aligned well with RsgI (**Table S1**). These findings prompted us to consider that RsgI’s JMD could have enzymatic activity. Specifically, we wondered whether this domain undergoes autoproteolysis. In reading about extracytoplasmic autoproteases, we noticed that the AlphaFold2 model of RsgI’s JMD had a similar fold to eukaryotic sea urchin sperm protein, enterokinase, agrin (SEA) domains (17, 23). SEA domains are broadly conserved among eukaryotes but homologs have not been identified in bacteria. Several of these domains have been shown to undergo autoproteolysis and remain noncovalently associated following cleavage. Furthermore, some have been implicated in mechanotransduction. Although SEA domains were not identified as structurally similar to RsgI’s JMD using the DALI server, both consist of alternating α- helices and ß-strands that form a ß-sheet packed at its concave face against two α-helices (**Fig. 1A-C**). A structural alignment using TM-align (24) of RsgI’s JMD with the predicted structure of the most well studied SEA domain, *Homo sapiens* Mucin-1 (MUC1), revealed a root-mean-square deviation (RMSD) of 4.82 Å over 75 amino acids, and a TM-align score of 0.35, further suggesting that the domains are unlikely to be homologous (**Fig. S2**). Furthermore, RsgI’s JMD and the MUC1 SEA domain contain similar secondary structural elements but their interconnectivities are distinct (**Fig. 1D**). Thus, the JMD and SEA domains likely evolved convergently to adopt a similar fold. Based on the experiments presented below, we have named RsgI’s JMD a SEAL domain for SEA-like.

### B. subtilis RsgI undergoes autoproteolysis

Many SEA domains undergo autoproteolysis at a conserved site in a ß-hairpin and remain noncovalently associated following cleavage (14, 15, 25) Although RsgI’s SEAL domain lacks the autoproteolytic site found in SEA domains, we investigated whether it undergoes autoproteolysis. We fused the soluble SEAL domain to an N-terminal His-SUMO tag and expressed it in *Escherichia coli* followed by Ni^2+^-affinity chromatography. The purification yielded three polypeptides of ∼35, 15, 14 kDas (**Fig. 2A**). Mass spectrometry revealed that the 35 kDa species was the full-length His-SUMO-SEAL fusion. The two smaller polypeptides corresponded to the N- and C-terminal fragments derived from cleavage within the SEAL domain. Importantly, incubation of the purified proteins at 37 °C resulted in further loss of the full-length protein and accumulation of the cleaved products (**Fig. 2B**), consistent with autocleavage. However, to exclude the possibility that a contaminating *E. coli* protease was responsible for the observed proteolysis, we performed protease-free in vitro transcription-translation reactions using ^35^S-methinonine and a *gfp- rsgI(SEAL)* fusion as a template. After a 60 min reaction, we observed full-length GFP-SEAL and two cleavage products (**Fig. 2C**). Inhibition of protein synthesis with chloramphenicol at 60 min resulted in further loss of the full-length species. Taken together, these data suggest that *B. subtilis* RsgI’s SEAL domain is an autoprotease.

**Figure 2.**
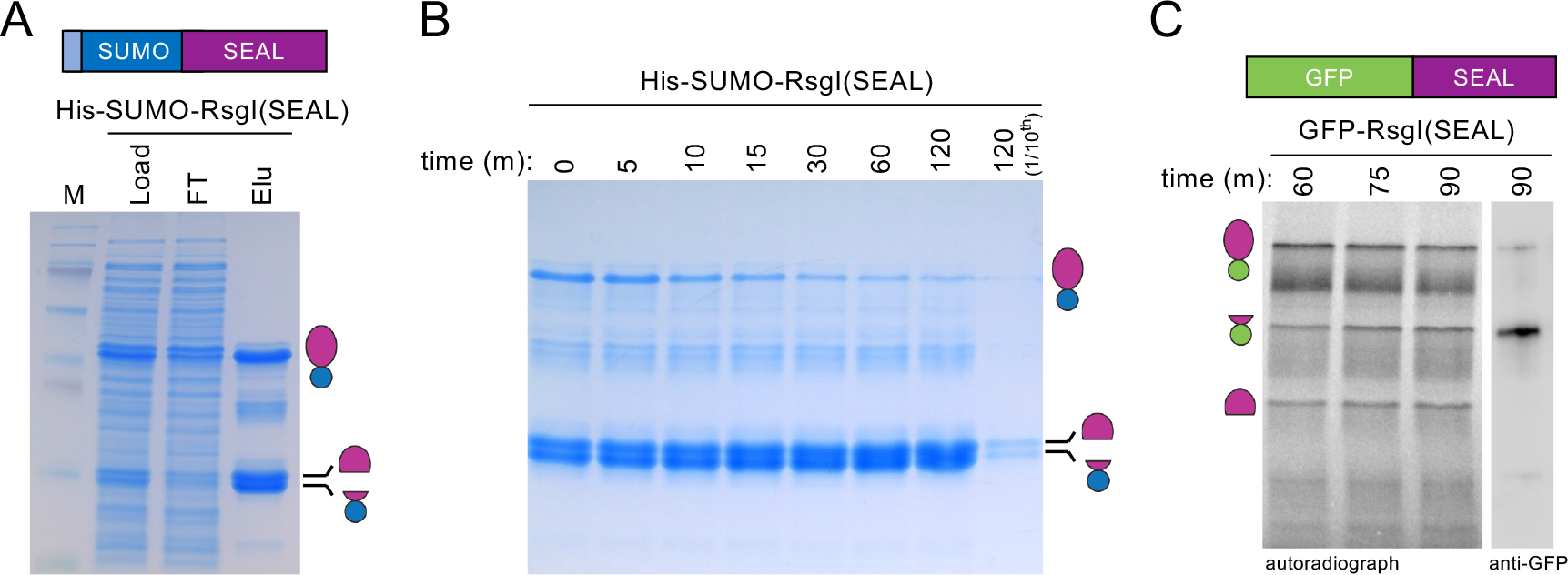
RsgI’s SEAL domain undergoes autoproteolysis. (A) Coomassie-stained SDS-PAGE gel of His-SUMO-RsgI(SEAL) expressed and purified from *E. coli.* His-SUMO-RsgI(SEAL) was expressed for 3 h, and the clarified lysate (Load) was subjected to affinity chromatography. The flowthrough (FT) was collected and His-SUMO-RsgI(SEAL) eluted (Elu) by the addition of imidazole. Full-length His-SUMO-RsgI(SEAL) and cleaved products are indicated. **(B)** Coomassie-stained gel of the elution in (A) at the indicated times (in min) after incubation at 37 °C. **(C)** Autoradiograph and immunoblot of an in vitro transcription-translation reaction with ^35^S-methionine using *gfp-rsgI(SEAL)* as template. Chloramphenicol was added 60 min after the reaction was initiated to inhibit protein synthesis. 75 and 90 min timepoints have an increase in cleavage products and a reduction in full- length protein. Anti-GFP immunoblot is from the 90 min reaction diluted 1:50. The purification, timecourse, and in vitro transcription-translation reaction were performed in biological triplicate and representative gels are shown.

### Autoproteolysis is a conserved feature among RsgI homologs

Many Clostridial species encode SigI/RsgI homologs and several of these have been found to regulate the expression of genes involved in the degradation of cellulose and other plant cell wall polysaccharides (26–29). The genes encoding cellulolytic enzymes are induced in the presence of these large polysaccharides that are unable to cross the bacterial cell wall. Several Clostridial RsgI homologs have carbohydrate binding modules (CBMs) appended to their IDRs and we previously proposed that cellulose binding to these domains generates a pulling force that could dissociate the site-1-cleaved products (**Fig. 3A**) (4). However, it was unknown whether these RsgIs are cleaved by a site-1 protease. Prompted by our discovery that *B. subtilis* RsgI is an autoprotease, we investigated whether Clostridial RsgI homologs also undergo autoproteolysis. We expressed and purified His-SUMO-fusions to the SEAL domains of RsgI from *Clostridium thermocaliphilum*, and RsgI2 and RsgI4 from *Hungateiclostridium thermocellum*. All three purifications contained cleavage products and a small amount of full-length protein (**Fig. 3B**). Similar to *B. subtilis* RsgI’s SEAL domain, the full-length *C. thermocaliphilum* His-SUMO-SEAL fusion was further lost during incubation at 37 °C (**Fig. 3C**). These data argue that the SEAL domains of Clostridial RsgI homologs undergo autoproteolysis and provide support for the model that plant polysaccharides trigger SigI activation via mechanotransduction (**Fig. 3A**).

**Figure 3.**
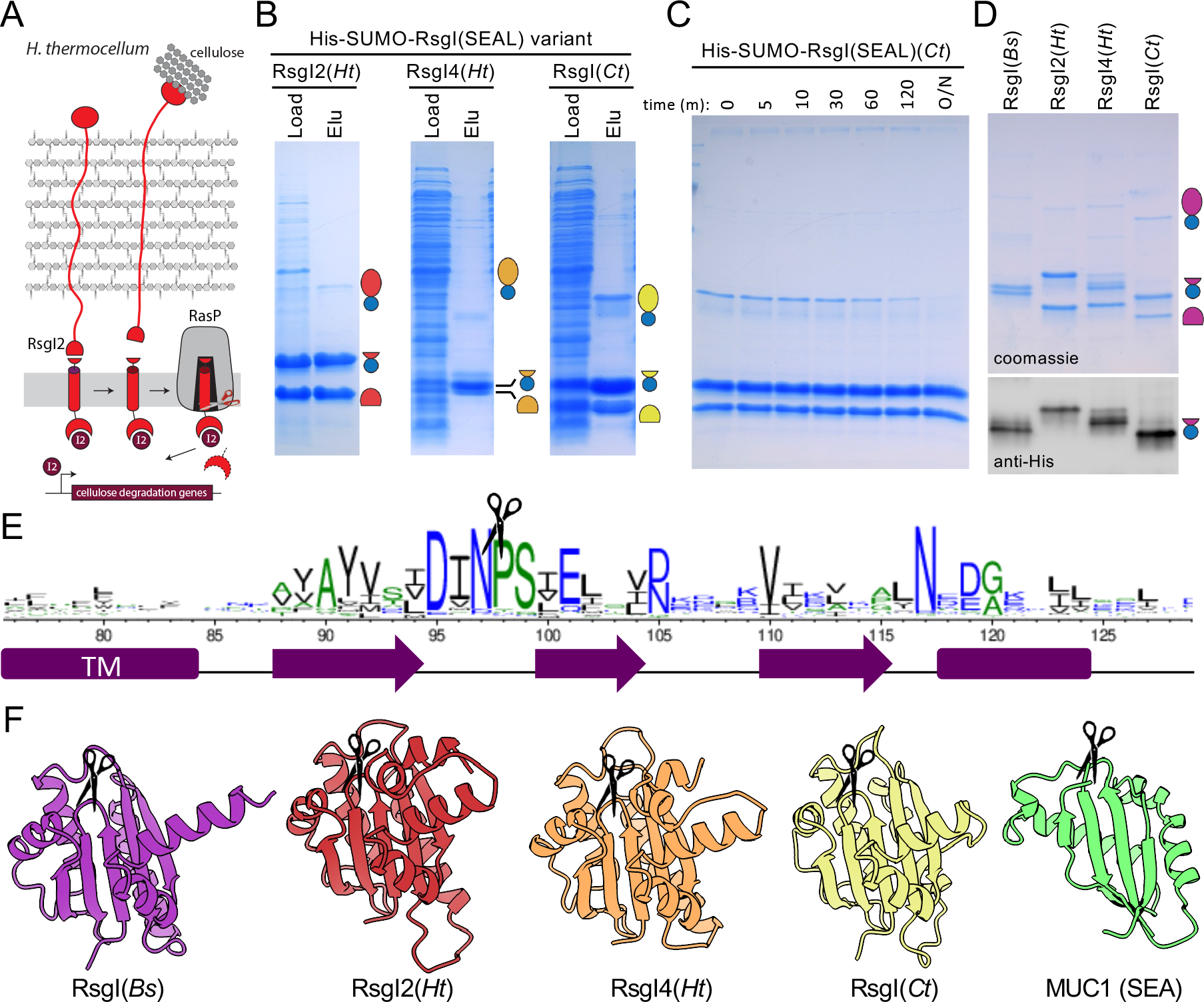
**RsgI homologs undergo autoproteolysis at a conserved cleavage site**. **(A)** Schematic model of the RsgI2-Sig2 signal transduction pathway in *H. thermocellum.* Cellulose binding to the carbohydrate binding module on RsgI2 generates a pulling force that dissociates the cleaved SEAL domain and enables RasP-mediated intramembrane proteolysis. Release of SigI2 (I2) activates genes involved in cellulose degradation. **(B)** Coomassie-stained gels of His- SUMO-RsgI(SEAL) fusions from *H. thermocellum* (*Ht*) RsgI2 and RsgI4 and *C. thermocaliphilum* (*Ct*) RsgI. Load and elution (Elu) are shown. All purifications were performed in biological triplicate and a representative gel is shown. **(C)** Coomassie-stained gel of purified His-SUMO- RsgI(SEAL)(*Ct*) after incubation at 37 °C for the indicated times. **(D)** Coomassie-stained gel and immunoblot of purified His-SUMO-RsgI(SEAL) variants. Purified proteins were diluted 1:50 for the immunoblot. **(E)** Sequence logo determined from the alignment of >5000 RsgI homologs. Adapted from *Brunet et al*. (4). The autocleavage site is highlighted (scissors). **(F)** AlphaFold2- predicted structures of SEAL domains from RsgI homologs with their cleavage sites (scissor) indicated. The crystal structure and cleavage site of MUC1’s SEA domain are included for comparison.

To identify the cleavage site within the SEAL domains, we performed Edman degradation on the C-terminal cleavage products (**Fig. 3D**). For all SEAL domains analyzed, the cleavage site mapped between N97 and P98 (**Fig. 3E, S3**). The cleavage site was independently confirmed by top-down (intact) mass spectrometry using MALDI-TOF (**Fig. S4**). Importantly, our previous mass spectrometry analysis of the in vivo cleavage products of *B. subtilis* RsgI was consistent with site-1 cleavage occurring at this exact position (**Fig. S5**). The cleavage site lies in a conserved 3 amino acid ß-hairpin between two ß-strands in the predicted structures (**Fig. 3F**). Importantly, cleavage is predicted to leave the ß-sheet intact and thereby allow ectodomain retention. This cleavage site is analogous to the cleavage site in the MUC1 SEA domain, which also lies in a ß-hairpin (15, 30), although there is no amino acid conservation between them (**Fig. 3F**).

### RsgI Autoproteolysis is catalyzed by conformational strain

Autoproteolysis of eukaryotic SEA domains is largely driven by conformational strain in the scissile ß- hairpin (15, 19, 20). In particular, amino acid substitutions of the nucleophilic residue in MUC1 impair but do not abrogate cleavage (31). Rather, insertion of glycine residues adjacent to the cleavage site to relax the torsional strain strongly impairs cleavage (23, 32). Our attempts to block autoproteolysis of RsgI’s SEAL domain by substituting conserved amino acids proximal to the cleavage site similarly slowed but did not abolish cleavage (**Fig. S6A**). Given the structural and functional similarities between the SEAL and SEA domains, we investigated whether relaxing the strain within the scissile loop of RsgI’s SEAL domain would abrogate cleavage. We inserted three glycine residues adjacent to the cleavage site (between I96 and N97) and purified the mutant from *E. coli* (**Fig. 4A**). Strikingly, purified His-SUMO-SEAL^GGG^ was almost entirely full-length (**Fig. 4B**). Furthermore, we detected no loss of the full-length product after incubation of the purified protein at 37 °C for >10 hours (**Fig. 4C, S6B**). Finally, a protease-free in vitro transcription-translation reaction using *gfp-rsgI(SEAL^GGG^)* as a template produced only the full-length product (**Fig. 4D**). Collectively, these data indicate that autoproteolysis of RsgI is catalyzed by conformational strain.

**Figure 4.**
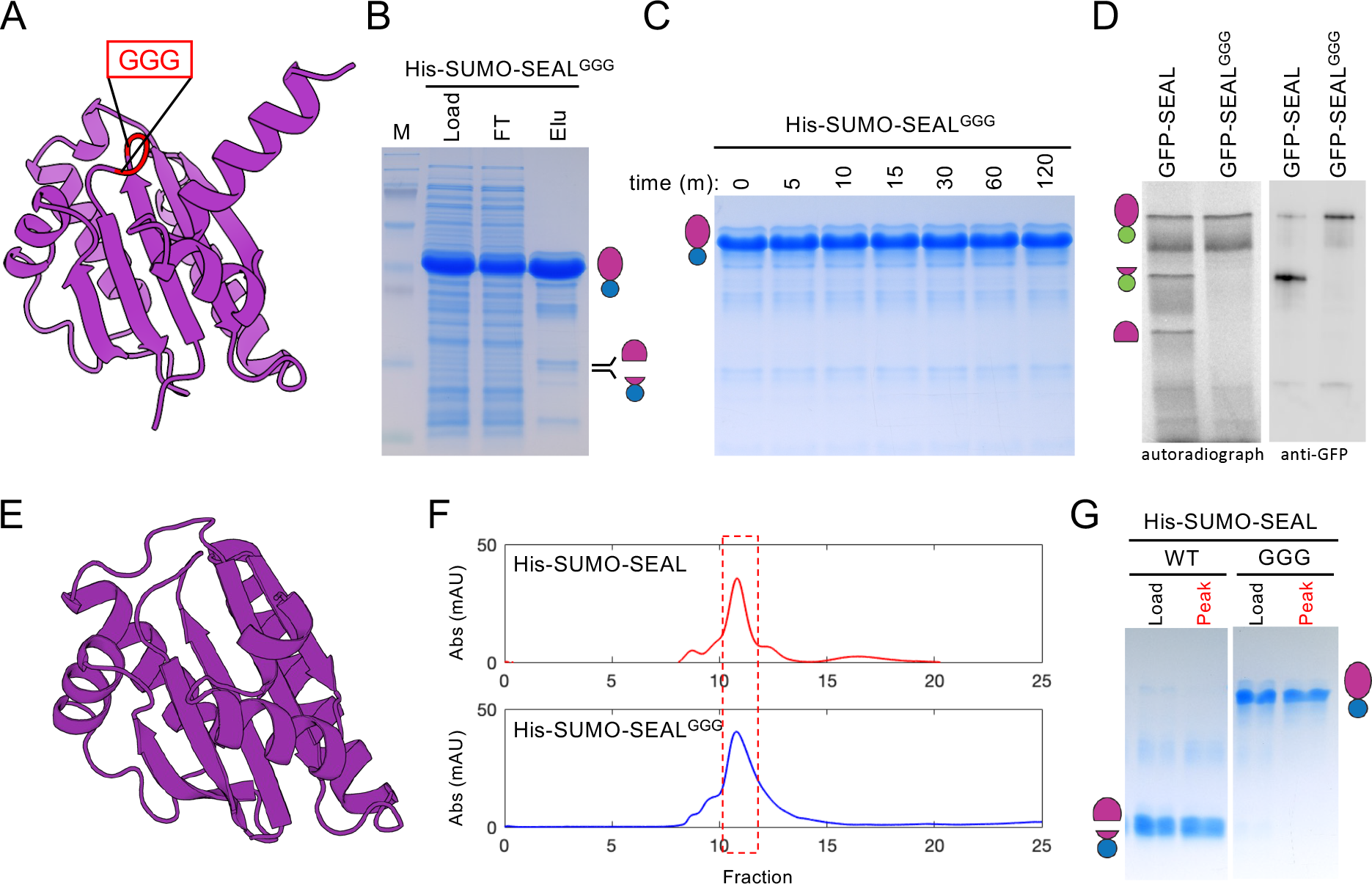
Autoproteolysis of RsgI’s SEAL domain is catalyzed by conformational strain. **(A)** AlphaFold2 model of SEAL^GGG^ with the inserted glycines in red. **(B)** Coomassie-stained gel of His-SUMO-SEAL^GGG^ expressed and purified from *E. coli*. Load, flowthrough (FT), and eluate (Elu) from Ni^2+^-affinity chromatography are shown. **(C)** Coomassie-stained gel of purified His- SUMO-SEAL^GGG^ after incubation at 37°C for the indicated times in min. **(D)** Autoradiograph and immunoblot showing GFP-SEAL and GGG variant after a 90 min in vitro transcription-translation reaction with ^35^S-methionine. Reactions were diluted 1:100 for visualization by immunoblot. **(E)** Cartoon rendering of the crystal structure of the *B. subtilis* SEAL^GGG^ domain. **(F)** Size-exclusion chromatograms for purified His-SUMO-SEAL and His-SUMO- SEAL^GGG^. **(G)** SDS-PAGE gel of the peak fraction (boxed in red) compared to load. The purification, timecourse, and in vitro transcription-translation reaction were performed in biological triplicate and representative gels are shown.

### Crystal structure of the SEAL^GGG^ domain

To experimentally determine the structure of the SEAL domain, we took advantage of the stability of the SEAL^GGG^ variant. The His-SUMO-SEAL^GGG^ fusion was affinity purified and, following removal of the His-SUMO tag, further purified by size-exclusion chromatography. Crystals were raised via hanging drop vapor diffusion and the structure of the SEAL^GGG^ domain was determined at 1.9 Å resolution (**Fig. 4E**). The structure revealed a fold consisting of five ß-strands that form a ß-sheet packed against two α-helices occurring in the order ß1-ß2-ß3-α1-α2-ß4-α3-ß5-α4-α5, with α4 and α5 only resolved on one of the two molecules in the asymmetric unit. The entire experimentally determined SEAL^GGG^ structure was nearly superimposable on the AlphaFold2 model with a RMSD of 1.08 Å and a TM-align score of 0.97 (**Fig. S7**). The electron density in the glycine loop region was weak, suggesting that the loop is highly flexible and may consist of a mixture of cleaved and uncleaved species (**Fig. S8**). For simplicity, we have modeled it as the uncleaved variant in Figure 4E.

Interestingly, ß1 and ß2, which surround the ß-hairpin cleavage site, had extended ß-strands in the crystal structure compared to the AlphaFold2 prediction of the wild-type SEAL domain (**Fig. S9**). Specifically, I96 and P101 of the ß-hairpin are incorporated into ß1 and ß2, respectively, and the inserted glycine residues largely populate the ß-turn (**Fig. S9**). Notably, both the AlphaFold2 prediction of the SEAL^GGG^ variant and the AlphaFold-multimer prediction of the cleavage products predict a similar ß-strand-extension and provide a mechanism of strain relief in the loop (**Fig. S9**). Previous studies on SEA domains have shown that the short ß-hairpin turn precludes proper folding in the absence of autoproteolysis (23). Our structure suggests that the mechanism of strain-catalyzed cleavage is likely to be conserved between SEA and SEAL domains.

### RsgI’s autoproteolytic products remain stably associated

In both the SEAL^GGG^ structure and AlphaFold-multimer prediction of the site-1 cleavage products, the two ß-strands bridged by the ß-hairpin are extended after cleavage creating additional hydrogen bond contacts between them. In the context of SEAL domain autoproteolysis, the extended ß-strands would likely further stabilize the interaction between the two cleavage products. We note that the Ni^2+^-affinity purification of the His-SUMO-SEAL fusions contained both cleavage products even though only one of the two contained a His tag. This strongly suggests that the cleaved products remain stably associated through the hydrogen bond network in the ß-sheet. However, it was formally possible that the full-length His-SUMO-SEAL fusion autoproteolyzed during elution and the two products do not interact. To distinguish between these models, we performed analytical size exclusion chromatography on purified His-SUMO-SEAL and the GGG variant. As can be seen in Figure 4F, the peak fractions and overall traces were similar. Importantly, the peak fraction from the SEAL domain contained the two cleavage products while SEAL^GGG^ was full- length (**Fig. 4G**). We conclude that RsgI remains noncovalently associated after autoproteolysis.

### RsgI^GGG^ abrogates site-1 cleavage in vivo

To investigate whether RsgI undergoes autoproteolysis in vivo, we generated the triple-glycine mutant in the full-length protein. To visualize the RsgI cleavage products, we used a previously characterized variant with GFP fused to the N-terminus and a His-tag at the C-terminus (4). Exponentially growing *B. subtilis* cells expressing GFP-RsgI-His or the GGG mutant were harvested before and at time points after inhibition of translation to monitor the fate of RsgI and its cleavage products. As reported previously, only a minor fraction of wild-type RsgI was full-length and both the N- and C-terminal site-1 cleavage products were readily detectable (4). The N-terminal site-2 cleaved product was also observed. Furthermore, the cleavage products were largely stable after translation inhibition. By contrast, most of the RsgI^GGG^ variant was full-length with a small amount of site-1 cleaved fragments, similar to what was observed in vitro (**Fig. 5A**). We conclude that RsgI undergoes autoproteolysis in vivo and is responsible for the constitutive site-1 cleavage in this RIP signaling pathway.

**Figure 5.**
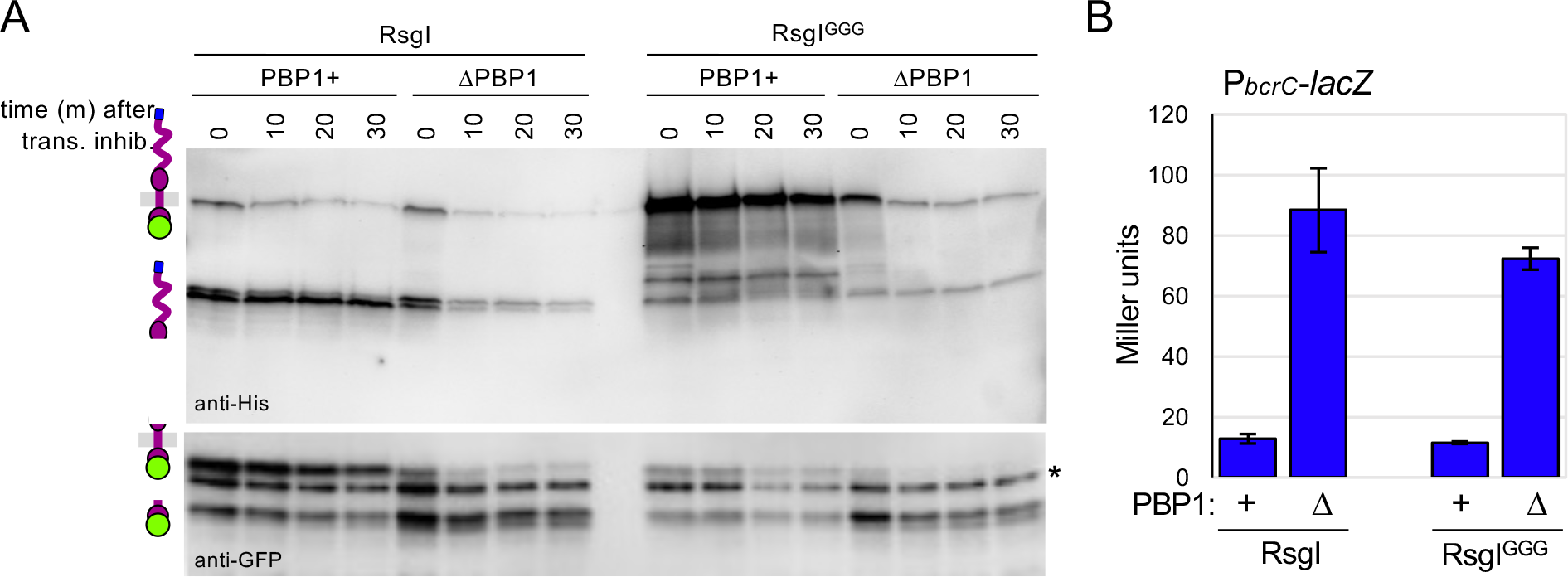
RsgI autoproteolysis occurs in vivo but is not required for SigI signaling. **(A)** Immunoblot of *B. subtilis* cells constitutively expressing GFP-RsgI-His or GFP-RsgI^GGG^-His in the indicated backgrounds. Timepoints (in minutes) before and after treatment with chloramphenicol and spectinomycin to inhibit protein synthesis are shown. Full-length GFP- RsgI^GGG^-His accumulates in cells with intact cell walls (PBP1+) but not in the ΔPBP1 mutant. A cross-reactive band in the anti-GFP immunoblot is indicated with an asterisk (*). The blot was performed in biological triplicate and a representative blot is shown. **(B)** Bar graph showing ß- galactosidase activity of a SigI-responsive (P*_bcrC_*) reporter. Strains harboring untagged RsgI and RsgI^GGG^ respond similarly to cell wall defects (ΔPBP1). All ß-galactosidase assays were performed in biological triplicate and error bars indicate standard error among these.

We have previously shown that the two site-1 cleavage products are rapidly lost in cells lacking the major cell wall synthase PBP1 (4). The N-terminal membrane fragment is cleaved by the membrane-embedded site-2 protease RasP leading to SigI activation, while the C-terminal fragment is degraded probably upon release into the medium. Importantly, SigI activation and loss of the two cleavage products does not occur if RsgI lacks its intrinsically disordered region (IDR) (4). These results led to the hypothesis that the IDR senses cell wall defects and generates a mechanical force that pulls the cleavage products apart, enabling intramembrane proteolysis and SigI activation. To investigate whether the RsgI^GGG^ variant is able to respond to cell wall stress, we analyzed the mutant in cells lacking PBP1. Surprisingly, the full-length RsgI^GGG^ protein was rapidly lost and the site-2 cleavage product was readily detectable. Furthermore, analysis of SigI activity using a SigI-responsive promoter (P*bcrC*) fused to *lacZ* (4) revealed that wild-type RsgI and the RsgI^GGG^ variant activate SigI similarly in the absence of PBP1 or after inhibition of its activity (**Fig. 5B and S10A**). These data indicate that intramembrane proteolysis and SigI activation can occur in response to cell wall defects independently of site-1 autoproteolysis.

### Intramembrane proteolysis in the absence of site-1 cleavage requires RsgI’s IDR

Our findings suggest that autoproteolysis occurs in vivo but preventing it is not sufficient to block intramembrane proteolysis and activation of SigI. One model that could explain these results is that the force required to dissociate the site-1-cleaved products is similar to the force required to partially or completely unfold the SEAL domain and once unfolded, it is cleaved by an unknown extracytoplasmic protease and/or becomes accessible to the membrane-embedded site-2 protease (**Fig. 6A**).

**Figure 6.**
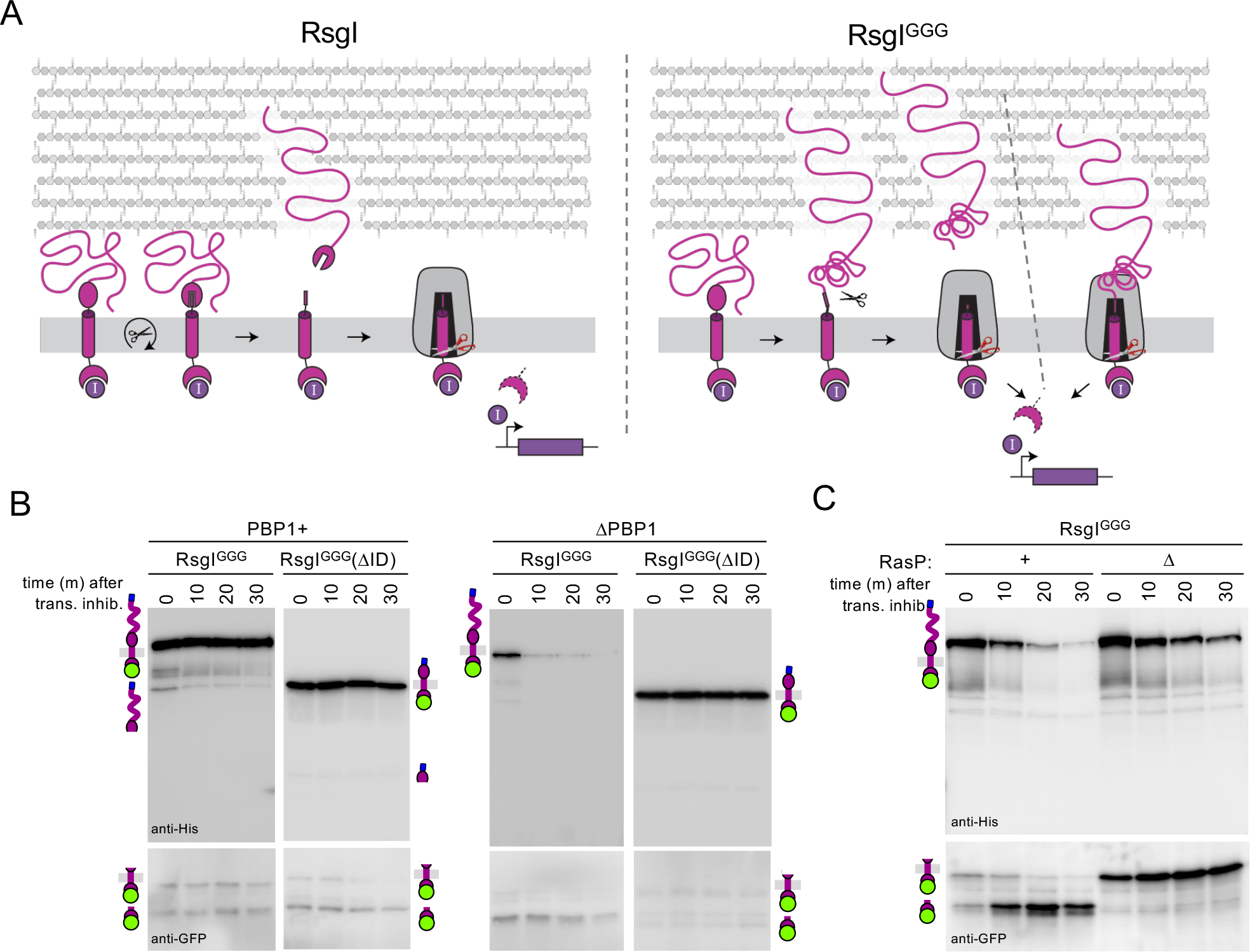
RsgI’s IDR is required to bypass site-1 autoproteolysis. (A) Schematic models of regulated intramembrane proteolysis of RsgI and the RsgI^GGG^ mutant. Detection of cell wall defects by RsgI’s IDR generates a force that pulls the autoproteolyzed SEAL domain apart enabling RasP-mediated intramembrane proteolysis (Left). The same pulling force unfolds the uncleaved SEAL^GGG^ domain, which becomes susceptible to an extracytoplasmic protease and/or can access the recessed interior of the RasP protease (Right). **(B)** Immunoblots of strains constitutively expressing the indicated GFP-RsgI^GGG^-His variants in the presence of abence of PBP1. Cell were harvested at the indicated times in minutes before and after inhibition of protein synthesis with chloramphenicol and spectinomycin. The intrinsically disordered region (ID) on RsgI^GGG^ is required to bypass autoproteolysis in the RIP signaling pathway. **(C)** Immunoblots of cell expressing GFP- RsgI^GGG^-His in the presence or absence of RasP. Cell were harvested at the indicated times in minutes before and after addition of moenomycin to inhibit PBP1 and chloramphenicol and spectinomycin to inhibit protein synthesis. All immunoblots were performed in biological triplicate and representative blots are shown.

Previous studies indicate that the IDR is required to dissociate the cleaved products enabling intramembrane proteolysis and SigI activation (4). To investigate whether the IDR was still required to trigger intramembrane proteolysis when site-1 cleavage was blocked, we analyzed the RsgI^GGG^ variant lacking its IDR. As can be seen in Figure 6B, the RsgI^GGG^ΔIDR mutant remained full-length in the presence *and* absence of PBP1. Similarly, activation of SigI is blocked in the RsgI^GGG^ΔIDR mutant (**Fig. S10B**). This suggests that the predicted mechanical force imparted by the IDR is required for signaling and that this force could drive unfolding of the SEAL domain.

To investigate whether RsgI^GGG^ is cleaved by an extracytoplasmic protease prior to intramembrane proteolysis or was directly processed by the site-2 protease RasP, we analyzed the mutant in cells lacking RasP. For these experiments, we used *rasP*^+^ and Δ*rasP* strains. Cells lacking RasP and PBP1 are not viable (4, 9). Accordingly, to generate cell wall defects to trigger signaling, we inhibited the glycosyltransferase activity of PBP1 using the drug moenomycin (33) prior to analyzing RsgI^GGG^. In the presence of the membrane-embedded protease RasP and moenomycin, full-length RsgI^GGG^ was efficiently cleaved by the site-2 protease similar to cells lacking PBP1 (**Fig. 6C**). In the absence of RasP, loss of the full-length product was markedly slower but a membrane-anchored RsgI fragment similar in size to the N-terminal fragment of autoproteolyzed RsgI accumulated. These data are consistent with a model in which partial or complete unfolding of the juxtamembrane domain in response to cell wall stress leads to cleavage by an extracytoplasmic protease that enables intramembrane proteolysis by RasP. This layered proteolysis is not unprecedented in mechanotransducive systems and is reminiscent of Notch signaling, in which a pulling force generates a conformational change in its ectodomain that enables cleavage by an extracellular protease to then trigger intramembrane proteolysis (34). However, we note that cleavage of RsgI^GGG^ by this unknown extracytoplasmic protease was slower in the absence of RasP, arguing that RasP is partially responsible for the direct processing of RsgI^GGG^ without prior proteolytic processing. Thus, our data suggest that partial or complete unfolding of the SEAL domain in response to cell wall defects enables cleavage by an extracytoplasmic site-1 protease but also allows RasP-mediated intramembrane proteolysis without ectodomain cleavage and release.

## DISCUSSION

Recent studies on the RsgI-SigI signaling system in *B. subtilis* revealed that RsgI is subject to regulated intramembrane proteolysis in response to cell wall defects (4, 9, 35). Analysis of the RsgI cleavage products in unperturbed cells strongly suggested that, unlike canonical RIP signaling pathways, site-1 cleavage is constitutive. The regulated step in this pathway was hypothesized to be the dissociation of the site-1 cleavage products by a mechanical force generated by RsgI’s intrinsically disordered region upon encountering cell wall defects. Here, we show that constitutive site-1 cleavage of RsgI is mediated by autoproteolysis and that the cleavage products remain stably associated, analogous to eukaryotic SEA domains. These data bolster the proposed model for RsgI-SigI signaling and provide support for the role of mechanical force in the activation of intramembrane proteolysis. Although the mechanism of force generation remains unclear in *B. subtilis*, our analysis of the Clostridial RsgIs suggests a clearly defined pulling force in the activation of their cognate sigma factors. Since cellulose and related carbohydrates are too large to enter the PG meshwork, it has been proposed that the IDRs on Clostridial RsgIs span the envelope layers displaying their carbohydrate binding modules on the cell surface (26, 27). Cellulose bound to these domains could generate a shear force that would dissociate the SEAL domain autocleavage products and trigger intramembrane proteolysis (**Fig. 3A**). Importantly, these Clostridial species encode RasP homologs, suggesting that their RsgI homologs are subject to RIP signaling. Future studies will focus on establishing whether Clostridial RsgIs are regulated by intramembrane proteolysis and determining how the IDR on *B. subtilis* RsgI generates a force when it encounters cell wall defects. Finally, these data lead us to propose that homologous SigI-RsgI signaling systems sense and respond to other extracellular macromolecules using the same mechanotransduction pathway.

### Bacterial SEAL domains

SEAL domains are broadly conserved among Firmicutes but largely absent in other bacterial phyla (**Fig. S11**). These domains are most often found in anti-sigma factors with a domain organization similar to *B. subtilis* RsgI. However, we identified many examples in which the anti-sigma factor had an additional domain appended to the IDR, as is found on *H. thermocellum* RsgIs (**Fig. S12**). In total, we found seven distinct domains appended onto the IDR of RsgI-like anti-sigma factors that contain SEAL domains (**Table S3**). In most cases, these domains are predicted to bind glycopolymers but others could bind extracellular proteins. In fact, the SEAL domain is frequently appended to PepSY domains, a poorly characterized domain that has been implicated in inhibiting extracellular enzymes (36). Interestingly, the SEAL-PepSY fusions lack an IDR and in some cases lack a cytoplasmic anti-sigma factor domain or even a TM segment. In virtually all cases, SEAL domain-containing proteins possess the highly conserved residues in the ß- hairpin involved in catalysis. Biochemical analysis of SEAL domains from two distinct protein families revealed that both undergo autoproteolysis in *E. coli* and the cleavage products noncovalently interact (**Fig. S13**). We hypothesize that most if not all SEAL domain-containing proteins autocleave yet remain associated. We further speculate that many of these proteins function in mechanotransduction.

### Common mechanisms of mechanotransduction

The structure of bacterial SEAL domains is similar to eukaryotic SEA domains, however, the interconnectivity of the structural elements are distinct, arguing that these domains arose from convergent evolution. It is therefore particularly striking how similarly these domains are employed in signal transduction. The most well studied SEA domain-containing protein, MUC1, highlights these similarities. MUC1 contains a cytoplasmic signaling domain, a single transmembrane helix, and an extracytoplasmic SEA domain followed by an extensive IDR (**Fig. 7**) (37). Both MUC1 and RsgI undergo constitutive autoproteolysis in their SEA/SEAL domains and their cleavage products remain noncovalently associated at the cell surface. Shear stress from pathogen binding to MUC1’s heavily glycosylated IDR has been implicated in dissociation of the N- and C-terminal fragments (38, 39). Although signaling through MUC1’s cytoplasmic domain is less well understood, evidence suggests that MUC1 is subject to intramembrane proteolysis by gamma-secretase (40). Furthermore, translocation of the MUC1’s free cytoplasmic domain to the cytoplasm, nucleus, and mitochondria has been documented (41). This signaling pathway requires further investigation but the work described here raises the exciting possibility that MUC1 could be regulated by a mechanotransduction pathway analogous to RsgI. Many other Mucins and cell surface proteins contain cytoplasmic signaling domains and extracytoplasmic SEA domains that undergo autoproteolysis suggesting that the these proteins could similarly function in RIP signaling.

**Figure 7.**
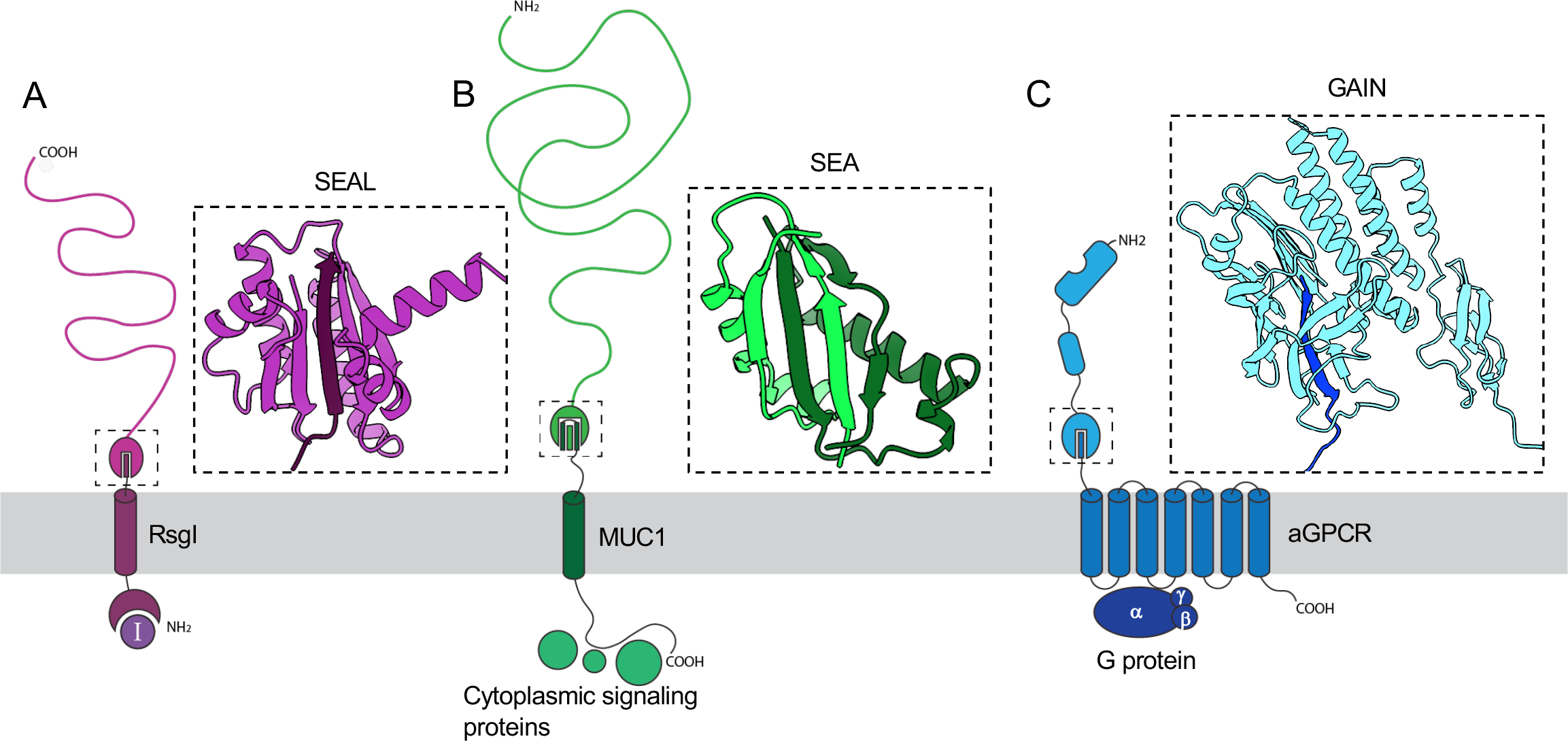
A common feature of mechanotransducive pathways. Schematic models of RsgI, Mucin-1 (MUC1), and an adhesion G Protein-Coulped Receptor (aGPCR). **(A)** RsgI undergoes autoproteolysis in its SEAL domain and the N (dark purple) and C-terminal (light purple) fragments remain non-covalently associated through an undisrupted ß-sheet. **(B)** MUC1 undergoes autoproteolysis in its SEA domain (pdb: 6bsb) and the C- (dark green) and N-terminal (light green) fragments remain noncovalently associated. Each cleaved product contributes two strands to the undisrupted ß-sheet. MUC1 has a large glycosylated extracellular intrinsically disordered region and cytoplasmic signaling proteins associate with its intracellular domain. **(C)** aGPCRs are autoproteolyzed within their GAIN domain (pdb: 4dlq), and the C-terminal (dark blue) and N-terminal (light blue) fragments remain noncovalently associated through an undisrupted ß-sheet. One or more extracellular adhesion domains are linked to the GAIN domain. The C-terminal strand of the GAIN domain is the tethered peptide agonist that, upon exposure, activates G-Protein signaling.

Unlike MUC1 signaling, adhesion GPCRs (aGPCRs) have been extensively characterized and the similarities to RsgI signaling merit discussion. aGPCRs undergo autoproteolysis in a highly conserved extracytoplasmic domain called the <u>G</u>PCR <u>A</u>utoproteolysis <u>IN</u>ducing (GAIN) domain (18). This domain does not resemble SEA or SEAL domains. However, the GAIN domain cleavage products remain stably associated through a ß-sheet network similar to SEA/SEAL domains (**Fig. 7**). Autoproteolysis occurs adjacent to the final ß-strand that connects to the GPCR transmembrane domain, much like the autocleavage site in RsgI. This ß-strand is termed the ‘tethered peptide’ and functions as an agonist of aGPCR signaling (42, 43). The peptide is shielded from interacting with the GPCR transmembrane domain when it is embedded in the ß-sheet network, but upon force imparted by the extracellular adhesion domains, the ectodomain is released exposing the peptide that, in turn, activates G Protein signaling (2). Interestingly, it has been shown that abolishing autocleavage in the GAIN domain is not sufficient to disrupt this signaling pathway (42, 44, 45). Recent structural data suggest that unfolding of the GAIN domain liberates the tethered peptide, which can still interact with the GPCR transmembrane domain without cleavage (45). Our data suggesting that unfolding the SEAL domain enables RasP-mediated intramembrane cleavage without ectodomain release shares striking parallels.

Our analysis of RsgI^GGG^ also indicates that an unidentified extracytoplasmic protease can cleave the partially or fully unfolded SEAL domain. This cleavage allows the membrane-anchored portion of RsgI to more efficiently access the recessed interior of the membrane-embedded protease. We suspect that this alternate site-1 cleavage is not normally used in *B. subtilis* when RsgI autoproteolysis is intact. However, we hypothesize that RsgI homologs exist that are not proficient in autoproteolysis but could still leverage mechanotransduction using site-1 proteases that cleave partially or fully unfolded SEAL domains. This type of RIP signaling would be analogous to the Notch pathway in which endocytosis generates a pulling force that unfolds the Negative Regulatory Region (NRR) of Notch, enabling site-1 cleavage by an ADAM family protease (46). We conclude by marveling at the strikingly similar mechanisms of mechanotransduction employed by bacteria and eukaryotes.

## METHODS

### General methods

All *B. subtilis* strains were derived from the prototrophic strain PY79 (47). Unless otherwise indicated, cells were grown in LB or defined rich (casein hydrolysate, CH) medium (48) at 37 °C. Insertion-deletion mutations were generated by isothermal assembly (49) of PCR products followed by direct transformation into *B. subtilis*. Tables of strains, plasmids and oligonucleotide primers and a description of strain and plasmid construction can be found online as supplementary data (Supplemental Table S4, S5, S6, S7, and Text S1).

### β-Galactosidase assays

*B. subtilis* strains were grown in LB medium at 37 °C to an OD600 of ∼0.7. The optical density was recorded and 1 mL of culture was harvested and assayed for β-galactosidase activity as previously described (50). Briefly, cell pellets were resuspended in 1 mL Z buffer (40 mM NaH2PO4, 60 mM Na2HPO4, 1 mM MgSO4, 10 mM KCl, and 50 mM β-mercaptoethanol). 250 µL of this suspension was added to 750 µL of Z buffer supplemented with lysozyme (0.25 mg/mL), and the samples were incubated at 37 °C for 15 min. The colorimetric reaction was initiated by addition of 200 µL of 2-nitrophenyl-β-D-galactopyranoside (ONPG, 4 mg/mL) in Z buffer and stopped with 500 µL 1M Na2CO3. The reaction time and the absorbance at 420 nm and OD550 of the reactions were recorded, and the β-galactosidase specific activity in Miller Units was calculated according to the formula [A420-1.75x(OD550)] / (time [min] x OD600) x dilution factor x 1,000 (51).

### Immunoblot analysis

Immunoblot analysis was performed as described previously (52). Briefly, 1 mL of culture was collected and resuspended in lysis buffer (20 mM Tris pH 7.0, 10 mM MgCl2 and 1mM EDTA, 1 mg/mL lysozyme, 10 µg/mL DNase I, 100 µg/mL RNase A, 1 mM PMSF) to a final OD600 of 10 for equivalent loading. The cells were incubated at 37 °C for 15 min followed by addition of an equal volume of Laemelli sample buffer (0.25 M Tris pH 6.8, 4% SDS, 20% glycerol, 10 mM EDTA) containing 10% β-mercaptoethanol. Samples were heated for 15 min at 65 °C prior to loading. Proteins were separated by SDS-PAGE on 12.5% polyacrylamide gels, electroblotted onto Immobilon-P membranes (Millipore) and blocked in 5% nonfat milk in phosphate-buffered saline (PBS) with 0.5% Tween-20. The blocked membranes were probed with anti-SigA (1:10,000) (53), anti-His (1:4,000) (GenScript), anti-GFP (1:10,000) (54) antibodies diluted into 3% BSA in PBS with 0.05% Tween-20. Primary antibodies were detected using horseradish peroxidase-conjugated goat anti-rabbit or anti-mouse IgG (BioRad) and the Super Signal chemiluminescence reagent as described by the manufacturer (Pierce). Signal was detected using a Bio-Techne FluorChem R System.

### *In vivo* protein turnover assay

In vivo protein turnover assays were performed as previously described (4). In brief, *B. subtilis* strains were grown in LB medium at 37 °C to an OD600 of 0.5. Protein translation was inhibited by the addition of both spectinomycin (200 µg/mL, final concentration) and chloramphenicol (10 µg/mL, final concentration). 1 mL samples normalized to an OD600 of 0.5 were collected immediately prior to antibiotic treatment and at the indicated times after. Cells were pelleted by centrifugation for 5 minutes and immediately flash-frozen in liquid nitrogen. The cell pellets were thawed on ice, lysed, and analyzed by immunoblot as described above.

### Purification of His-Sumo-RsgI and variants

Expression plasmids containing His-SUMO-RsgI variants were transformed into *E.coli* BL21(DE3) Δ*tonA* cells. Transformants were sub-cultured in terrific broth (TB) + 100 µg/mL ampicillin for 3 hours then back-diluted into 1 L TB + 100 µg/mL ampicillin to an OD600 of 0.01 and grown at 37 °C to an OD600 of 0.4. Cultures were induced with 500 µM IPTG for 3 hours at 37 °C and then harvested by centrifugation at 4000 rpm for 15 minutes. Pellets were resuspended in Lysis Buffer (50 mM HEPES-NaOH, 300 mM NaCl, 25 mM Imidazole, 10% (v/v) glycerol) and frozen at -80 °C. Cell pellets were thawed on ice and lysed by 2 passes through a cell disrupter at 25 kPsi. 125 U Benzonase (Sigma) and 1X complete protease inhibitor (Roche) were added to the cell lysate and incubated on ice for 15 min. Lysates were clarified by ultracentrifugation at 40,000 rpm for 45 minutes at 4 °C. The supernatant was passed two times over a column with 500 µL of Ni^2+^-NTA resin, washed with 50 bed volumes of wash buffer (20 mM HEPES-NaOH, 300 mM NaCl, 25 mM Imidazole, 10% (v/v) glycerol), and eluted in 5 mL of elution buffer (20 mM HEPES-NaOH, 300 mM NaCl, 300 mM Imidazole, 10% (v/v) glycerol).

When monitoring purified His-SUMO-RsgI variants for autoproteolysis, the protein concentration of each eluate was determined using a non-interfering protein concentration determination kit (G-Biosciences), and each prep was normalized to 1 mg/mL. Purified protein was placed at 37 °C and at indicated times points a sample was removed and mixed with an equal volume Laemelli SDS sample buffer containing 10% β-mercaptoethanol to stop the reaction. Samples were visualized by SDS-PAGE using 17.5% polyacrylamide gels and stained with Instant Blue (Abcam).

### Size-exclusion chromatography

His-SUMO-SEAL and His-SUMO-SEAL^GGG^ were analyzed by size-exclusion chromatography (SEC) on a Superdex S75 column (GE Healthcare) in buffer containing 20 mM of HEPES-NaOH, pH 7.5 and 300 mM of NaCl. The load and peak absorbance (A280) fractions, were diluted into equal volume 2X sample buffer containing 10% β-mercaptoethanol and resolved by SDS-PAGE using a 17.5% polyacrylamide gel and stained with Instant Blue (Abcam). Absorbance profiles were plotted using MATLAB.

### Edman Degradation and MALDI-TOF Analyses

Following purification of His-SUMO-RsgI homologs, the eluates were treated with 6.25 µg of Ulp1 and dialyzed at 4 °C overnight in Dialysis Buffer (20 mM HEPES-NaOH, 300 mM NaCl, 10% glycerol) with a 3kDa MWCO. The sample was loaded onto a BioRad column containing 500 µL of Ni-NTA resin and passed 3 times over the resin before the flowthrough containing purified RsgI variant was collected. The buffer was exchanged into water using a 3 kDa ultra-centrifugal filter (Sigma) and the protein analyzed by MALDI-TOF using an Applied Biosystems Voyager DE Pro in linear mode (Tufts University Core Facility).

Purified SEAL variants as described above were separated by SDS-PAGE using 17.5% polyacrylamide gels and transferred to a PVDF membrane at 90 V for 60 min. The membrane was then washed 5X with ddH2O, stained with 0.02% Coomassie Brilliant blue in 40% methanol, 5% acetic acid for 30 sec, destained in 40% methanol, 5% acetic acid for 1 min, washed 3X with ddH2O and allowed to air dry. The bands of interest were cut from the membrane and sent for 5 cycles of Edman Degradation analysis with an ABI 494 Protein Sequencer (Tufts University Core Facility).

### In vitro transcription-translation reactions

In vitro transcription-translation reactions were carried out using the NEB PURExpress system and its published protocol (https://www.neb.com/protocols/0001/01/01/protein-synthesis-reaction-using-purexpress-e6800). Briefly, Solution A, Solution B, and ^35^S-methionine were combined with 250 ng pAB87, pAB101, or pDHFR as a positive control. The reactions were incubated for 1 hour at 37 °C and 5 µL of the reaction was combined with equal volume Laemalli Sample Buffer containing 10% β-mercaptoethanol to stop the reaction. Chloramphenicol (25 µg/mL final) was added to the transcript-translation reaction after 60 min, to inhibit protein synthesis and samples were mixed with Laemelli Sample buffer at the indicated times. Proteins were separated by SDS-PAGE using a 17.5% polyacrylamide gel, the gel was dried, exposed to a phosphor screen, and imaged with an Amersham Typhoon IP (Cytiva). Samples were also visualized by immunoblot using anti-GFP antibodies as described above.

### RsgI^GGG^ purification for crystallography

pAB188 [His-Sumo-SEAL^GGG^] was transformed into BL21(DE3) Δ*tonA* cells. Transformants were subcultured in terrific broth (TB) with 100 µg/mL ampicillin at 37 °C for 3 hours then back-diluted into 3 L TB + 100 µg/mL ampicillin to an OD600 of 0.01 and grown at 37 °C to an OD600 of 0.4. Cultures were induced with 500 µM IPTG for 3 hours before collection by centrifugation at 4000 rpm. Pellets were resuspended in Lysis Buffer (50 mM HEPES-KOH pH 7.5, 250 mM NaCl, 25 mM Imidazole, 10% v/v glycerol) and frozen at -80 °C. Cell pellets were thawed on ice and lysed by two passes through a cell disrupter at 25k Psi. 125 U Benzonase (Sigma) and 1X complete protease inhibitor (Roche) were added to the cell lysate and incubated on ice for 15 min. Lysates were clarified by ultracentrifugation at 40,000 rpm for 45 minutes at 4 °C. The supernatant was passed two times over a column with 1.5 mL of Ni^2+^-NTA resin (Qiagen), washed with 50 bed volumes of wash buffer (20 mM HEPES-KOH, 250 mM NaCl, 25 mM Imidazole, 10% (v/v) glycerol), and eluted in 10 mL of elution buffer (20 mM HEPES-KOH, 250 mM NaCl, 300 mM Imidazole, 10% (v/v) glycerol). 6.25 µg of Ulp1 was added to the eluate to remove the His-SUMO tag and was dialyzed (3 kDa MWCO) overnight at 4 °C in dialysis buffer (20 mM HEPES-KOH, 250 mM NaCl, 10% v/v glycerol, 1 mM TCEP). The dialyzed elution was passed over a column containing 1.5 mL of Ni^2+^-NTA resin three times and the flowthrough collected and analyzed for purity by SDS-PAGE. The SEAL^GGG^ domain was purified by size exclusion chromatography (SEC) on a Superdex S75 column (GE Healthcare) in buffer containing 20 mM of HEPES-KOH pH 7.5, 250 mM of NaCl, and 1 mM TCEP. The peak absorbance (A280) fraction was collected and concentrated to 22 mg/mL using a 3 kDa MWCO concentrator (Millipore) before flash freezing in liquid nitrogen.

### Crystallization and structure determination

RsgI^GGG^ was crystallized using the hanging drop vapor diffusion method at 18 °C. Native protein was thawed on ice and diluted into a buffer solution (20 mM HEPES-KOH pH 7.5, 20 mM NaCl, 1 mM TCEP) to a final concentration of 8 mg/mL. Crystals were grown by mixing 1 µL of purified protein with 1 µL of a reservoir solution of 100 mM HEPES-KOH pH 7.5, 100 mM ammonium acetate, and 28% PEG-3350 (w/v) in EasyXtal 15-well trays (NeXtal) containing 400 µL reservoir solution. RsgI^GGG^ crystallized over 7 days and crystals were harvested and cryoprotected in reservoir solution supplemented with 15% ethylene glycol before freezing in liquid nitrogen.

X-ray diffraction data were collected at the Advanced Photon Source (beamline 24-ID-C) and data were processed using the SSRL autoxds script (A. Gonzalez, Stanford SSRL). An AlphaFold2 model of RsgI^GGG^ was used for molecular replacement to determine phase information and an initial map was determined using the Phaser program in Phenix v1.20.1 (55). Statistics were analyzed as described in Table S2. Model building was performed in Coot (56) and refinement was performed using Phenix. The structures are deposited in the PDB under code 8T9N.

### Bioinformatics Analysis

A local pblast run was performed on the amino acid sequence of *B.subtilis* RsgI’s SEAL domain against the RefSeq select database using an e-value cutoff of 0.05. The resulting scientific names were converted to taxids using NCBI’s taxid identifier (https://www.ncbi.nlm.nih.gov/Taxonomy/TaxIdentifier/tax_ identifier.cgi) (57) and taxids were used to plot onto a phylogenetic tree of 5767 unique bacterial taxa. The tree was constructed using iTOL (https://itol.embl.de/) (58).

The resulting FASTA files from the pblast search were annotated using local PfamScan (59) (https://www.ebi.ac.uk/seqdb/confluence/display/THD/PfamScan) against the Pfam database. The output domains were organized based on Refseq accession number and the resulting domain organizations built. Counts of each domains identified can be found in supplementary data (Table S3).

### Multiple Sequence Alignment

Multiple sequence alignments were generated using Clustal Omega (https://www.ebi.ac.uk/Tools/msa/clustalo/) (59).

### Alphafold2 Predictions

Protein structures were modeled using AlphaFold2 (21) and ColabFold run using the Alphafold2 Advanced Colab notebook (60) (https://colab.research.google.com/github/sokrypton/ColabFold/blob/main/beta/ AlphaFold2_advanced.ipynb) or downloaded (A0A6M4JFI2, A3DC27, A3DCG3, A0A1V4I8Y9) from the Alphafold database (61) (available at: https://alphafold.ebi.ac.uk/). The parameters for the run were as follows: msa method mmseq2, pair mode unpaired, maximum recycling 3, no templates, five models generated. The highest ranked structure by pLDDT is shown.

### Structural model visualization

Crystal structures of MUC1 (pdb: 6bsb) and aGPCR (pdb: 4dlq) were downloaded from the PDB. ChimeraX1.3 and Pymol 2.4.0 were used to visualize the structural models and generate images. PDBsum was used to generate MUC1 SEA and RsgI SEAL protein topology maps (62).

## Acknowledgements

We thank all members of the Bernhardt-Rudner supergroup for helpful advice, discussions, and encouragement, Kailey Slavik, Yao Li, and the Kranzusch lab for help with in vitro transcription-translation reactions, crystallization trials, and structure determination, Lior Artzi for Clostridial genomic DNAs, Suzanne Walker for moenomycin A, the Tufts University Core Facility (TUCF) and Taplin Biological Mass Spectrometry Facility for proteomic analysis, and Ernst Schmid for help with analysis of Alphafold2 predictions. Portions of this research were conducted on the O2 High Performance Computing Cluster, which is supported by the Research Computing Group at Harvard Medical School. Support for this work comes from the National Institute of Health Grants R35GM145299, U19 AI158028 (D.Z.R.). A.P.B. was funded in part by NSF (DGE1745303).

## Contributions

A.P.B. and D.Z.R conceived the study, A.P.B. performed the experiments and analyses, C.H. provided reagents, S.J.H. and P.J.K assisted with structure determination, D.Z.R supervised the study, A.P.B. and D.Z.R wrote the paper with edits from C.H., S.J.H., and P.J.K.

## Competing interests

The authors declare that they have no competing interests.

## Data availability

Raw data for all graphs has been uploaded as source data. Uncropped immunoblots and SDS-PAGE gels are included in the supplementary information. Plasmids, primers, synthetic DNA constructs and strains used can be found in supplementary tables. The X-ray crystal structure of the SEAL^GGG^ domain has been deposited in the Protein Data Bank (available at: https://www.rcsb.org/) with accession code 8T9N.

---

**Figure S1.**
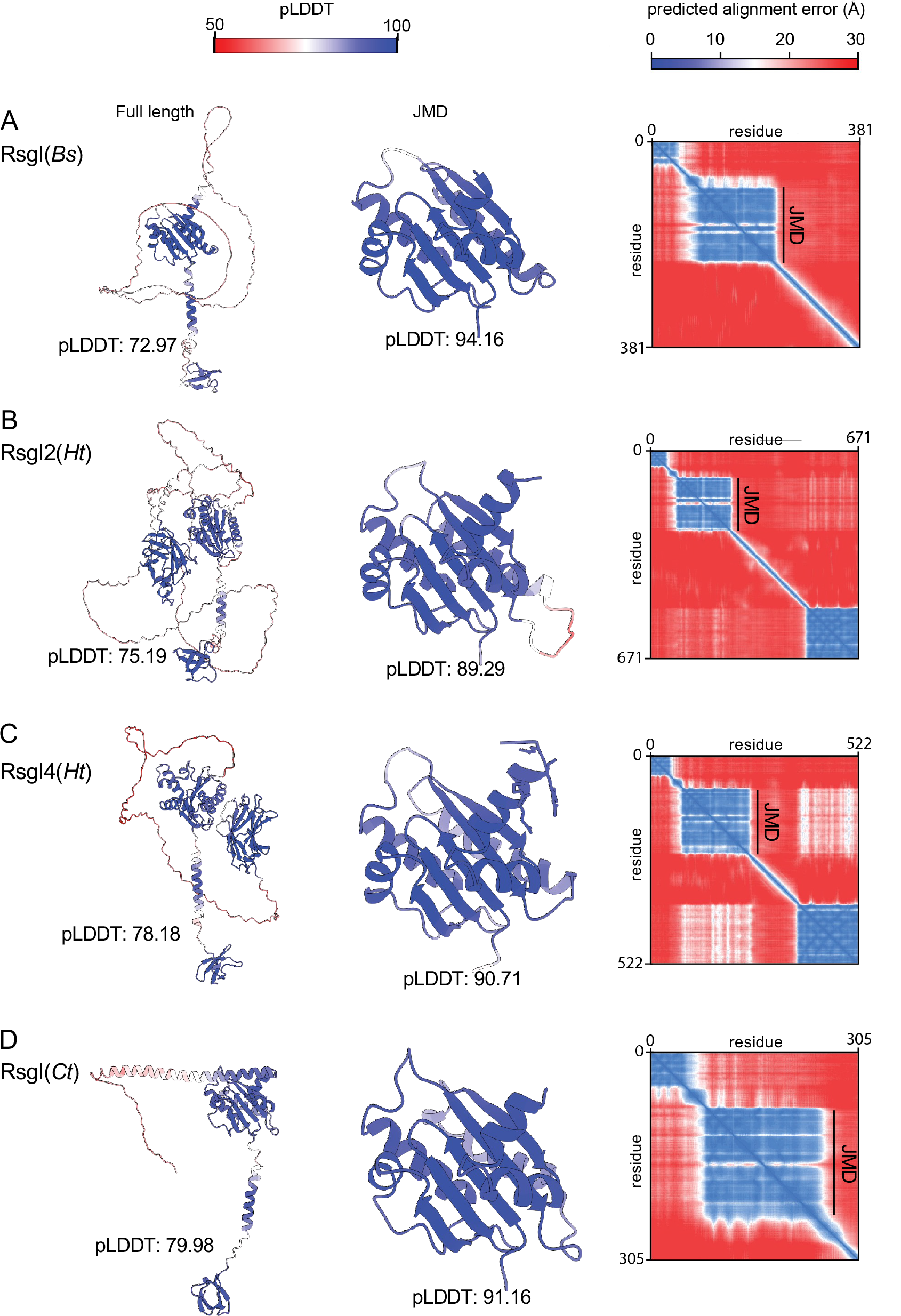
AlphaFold2 confidently predicts the juxtamembrane domain of RsgI homologs. AlphaFold-predicted structures colored by pLDDT, and pAE plots of **(A)** *B. subtilis* (*Bs*) RsgI; **(B)** *H. thermocellum* (*Ht*) RsgI2; **(C)** *H. thermocellum* RsgI4; and **(D)** *C. thermocaliphilum (Ct)* RsgI. The juxtamembrane domain (JMD) is highlighted on the pAE plots.

**Figure S2.**
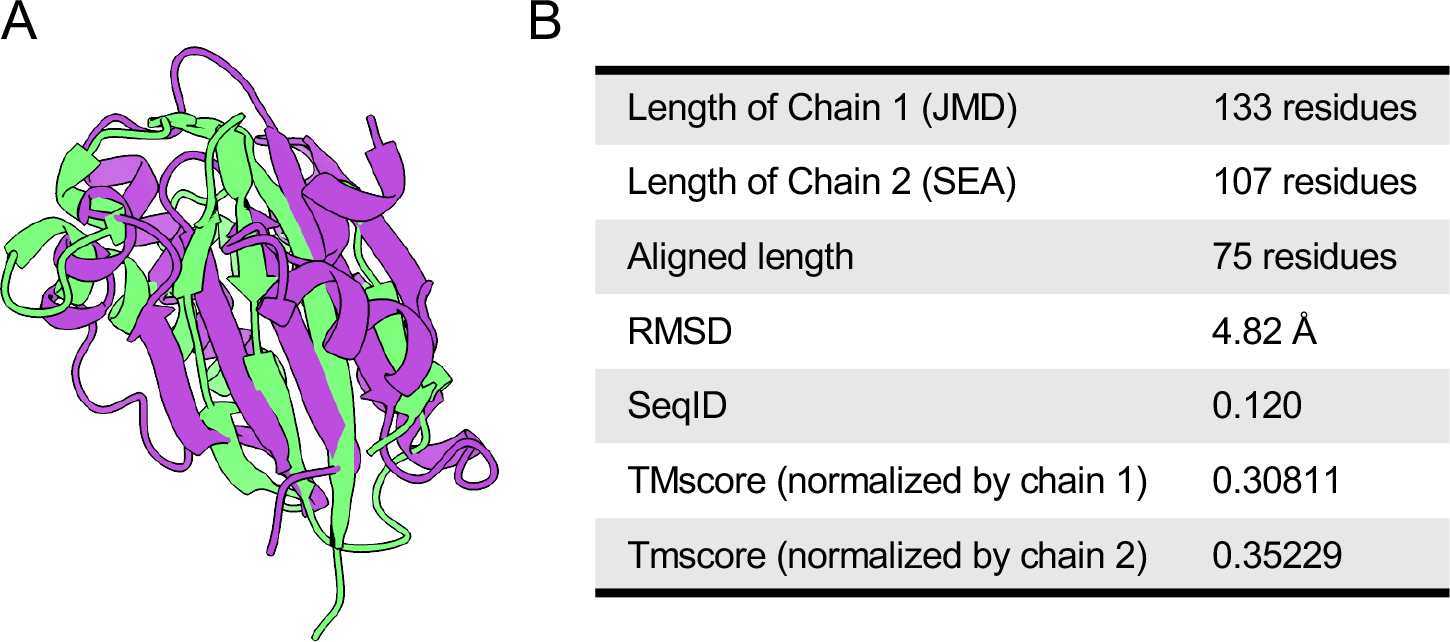
RsgI’s JMD and the SEA domain of MUC1 align by TM-align. **(A)** structural alignment of RsgI’s JMD (purple) and MUC1’s SEA domain (green). **(B)** Metrics for the domains and their alignment.

**Figure S3.**
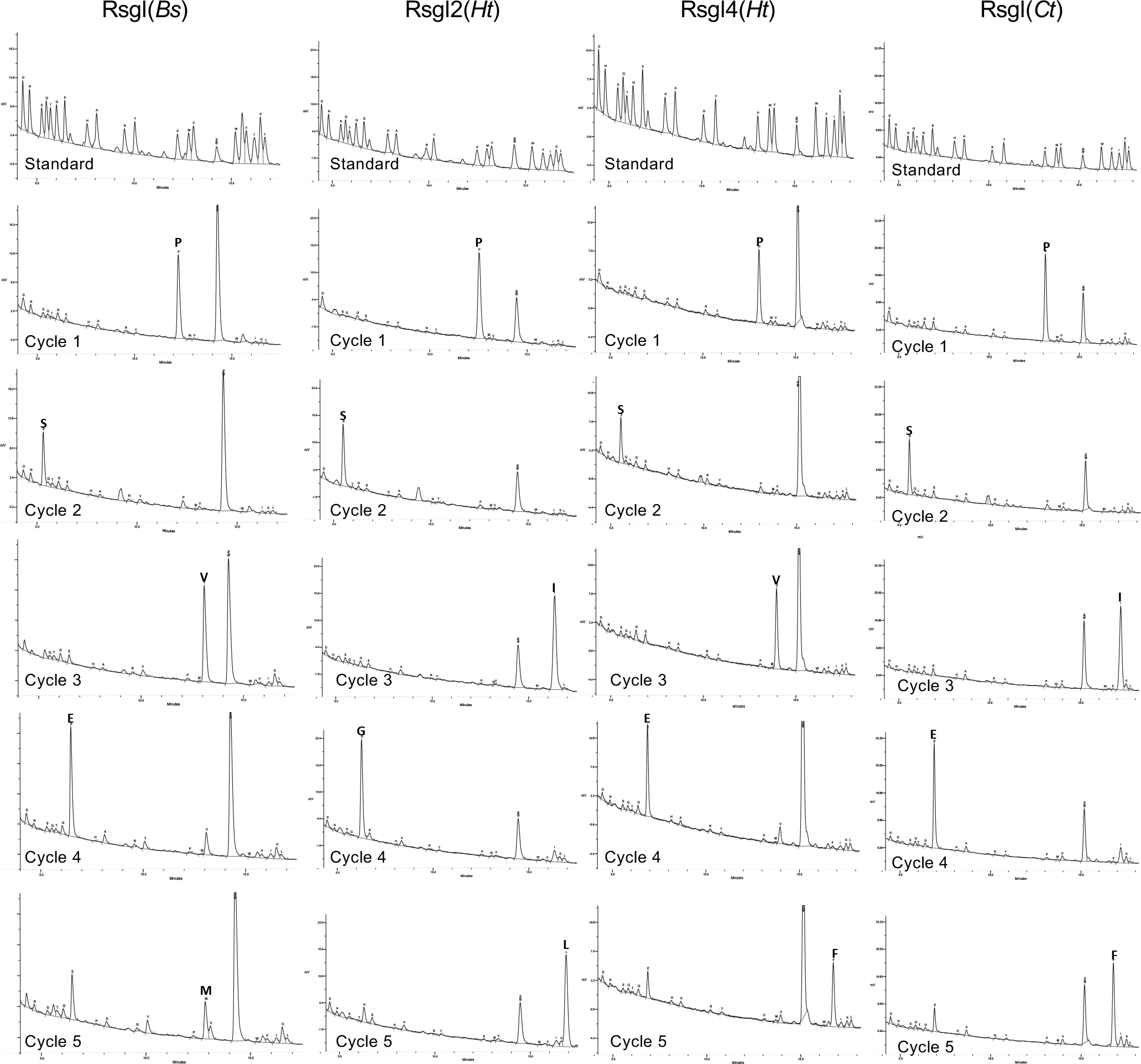
Edman degradation reveals a conserved cleavage site. Chromatograms from 5 cycles of Edman degradation for each of the four RsgI homologs tested as compared to standards of each amino acid. SEAL domains from each homolog were expressed and purified from *E. coli* and the N-terminal peptide sequence was determined from the C-terminal cleavage product.

**Figure S4.**
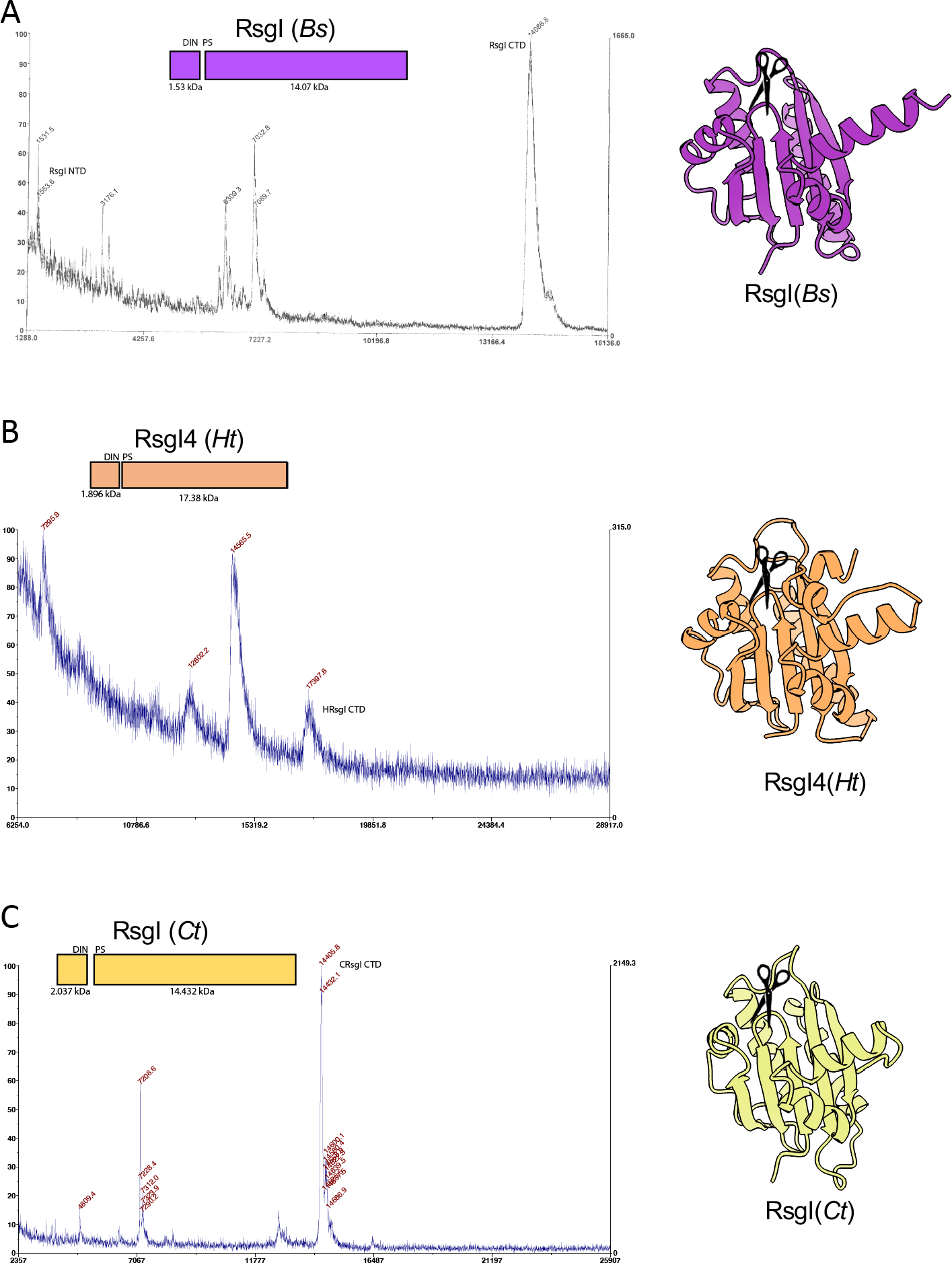
Intact mass spectrometry reveals a conserved cleavage site for RsgI homologs. Mass spectra of purified RsgI(SEAL) variants lacking His-SUMO tags. Protein diagrams indicating the predicted masses of the cleavage products are shown above each spectrum. AlphaFold-predicted structures and cleavage sites (scissors) are shown on the right.

**Figure S5.**
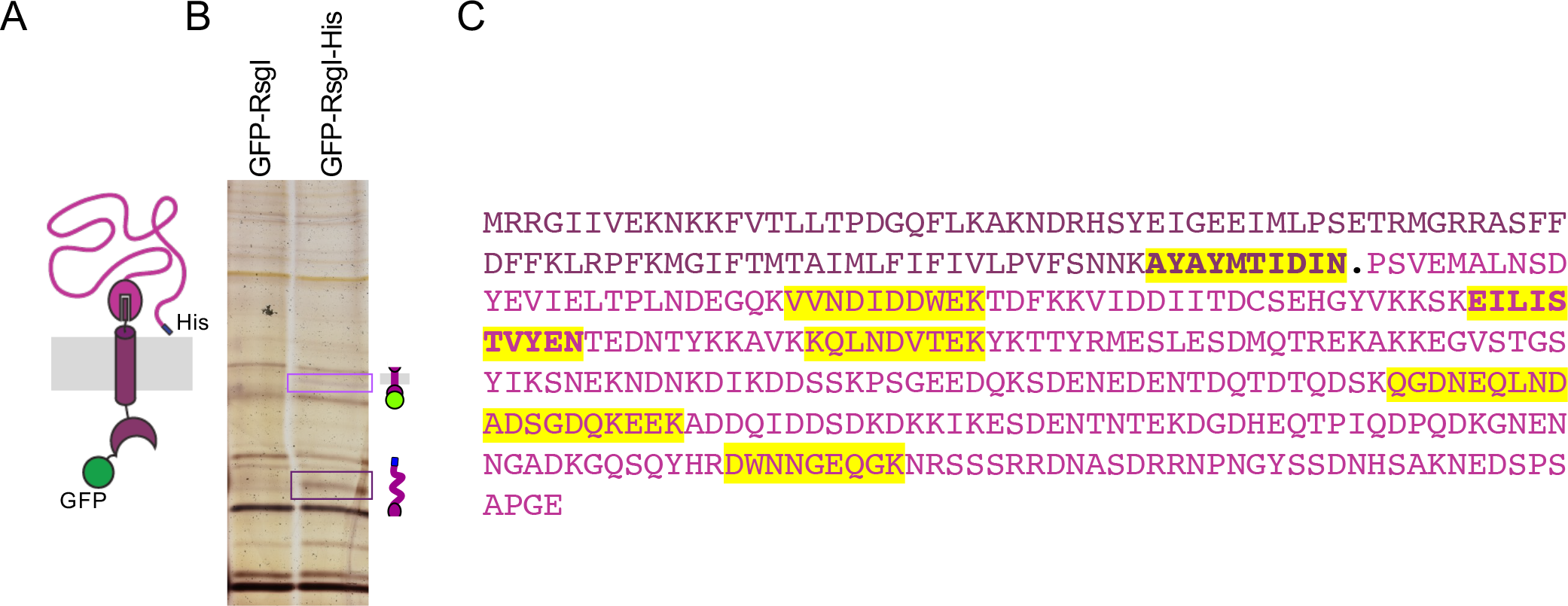
Site-1 cleavage in vivo is consistent with the autocleavage site in vitro. **(A)** Schematic of the GFP-RsgI-His6 fusion used in the assay. **(B)** Silver-stained SDS-PAGE gel of eluates from Ni^2+^-affinity chromatography of detergent-solubilized membrane fractions from *B. subtilis* cells expressing GFP-RsgI-His6 or GFP-RsgI. The RsgI fragments (boxed in light and dark purple) were excised, digested with trypsin, and analyzed by mass spectrometry. **(C)** Sequence of RsgI with N- and C-terminal cleavage products based on the in vitro autocleavage analysis highlighted in different shades of purple. The period (.) indicates the autocleavage site identified in vitro. Sequences highlighted in yellow indicate peptides detected by MS. The two peptides in bold lack a canonical tryptic cleavage site and are therefore candidates for the site of site-1 cleavage.

**Figure S6.**
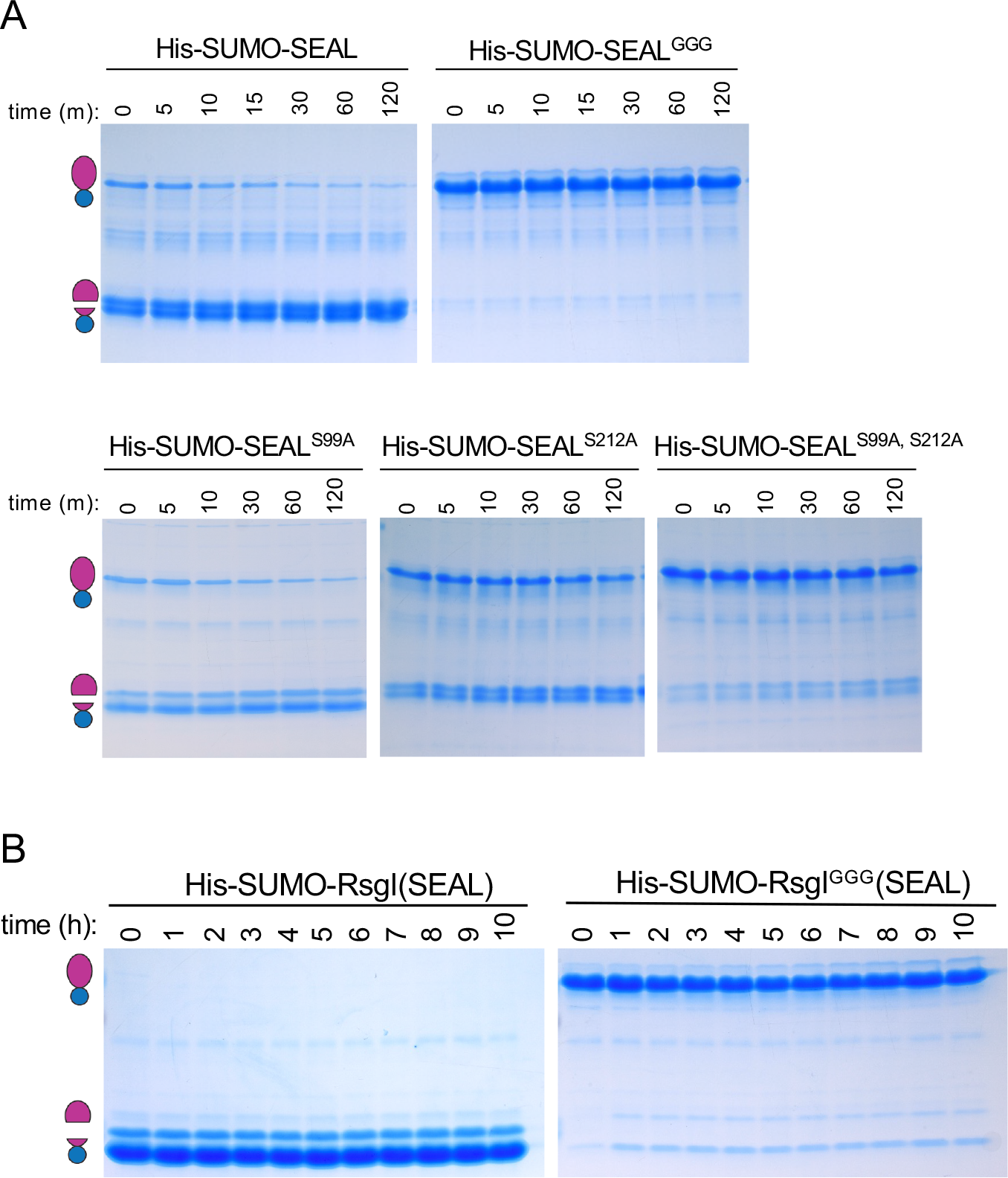
Point mutations in the SEAL domain fail to abolish autoproteolysis in vitro. **(A)** Commassie-stained gels of purified His-SUMO-SEAL and the indicated mutants incubated at 37 °C for the indicated time in minutes. All point mutants retain some autocleavage activity. **(B)** Coomassie-stained gels of purified His-SUMO-RsgI(SEAL) and the GGG variant incubated at 37 °C for the indicated time in hours. The SEAL^GGG^ variant undergoes modest autoproteolysis. All purifications and timecourses were performed at least two times.

**Figure S7.**
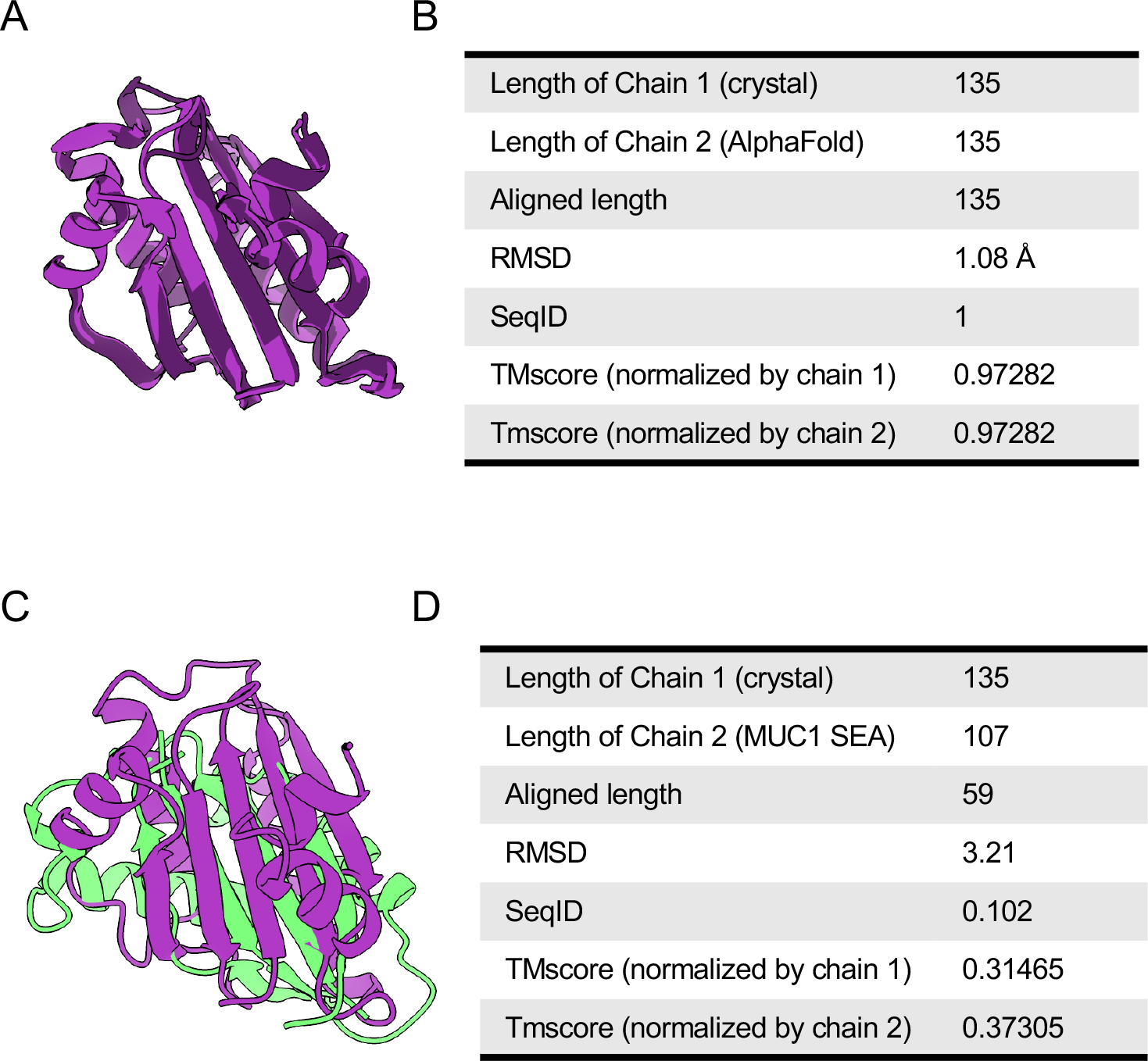
The X-ray crystal structure of SEAL^GGG^ closely resembles the AlphaFold2- predicted model and is structurally similar to MUC1’s SEA domain. (A) Structural alignment of the AlphaFold-predicted (dark purple) and experimentally determined (light purple) SEAL^GGG^ domains by TM-align. **(B)** Metrics describing the structural alignment. **(C)** Structural alignment of the x-ray crystal structure of SEAL^GGG^ and MUC1’s SEA (green) domains. **(D)** Metrics describing the structural alignment.

**Figure S8.**
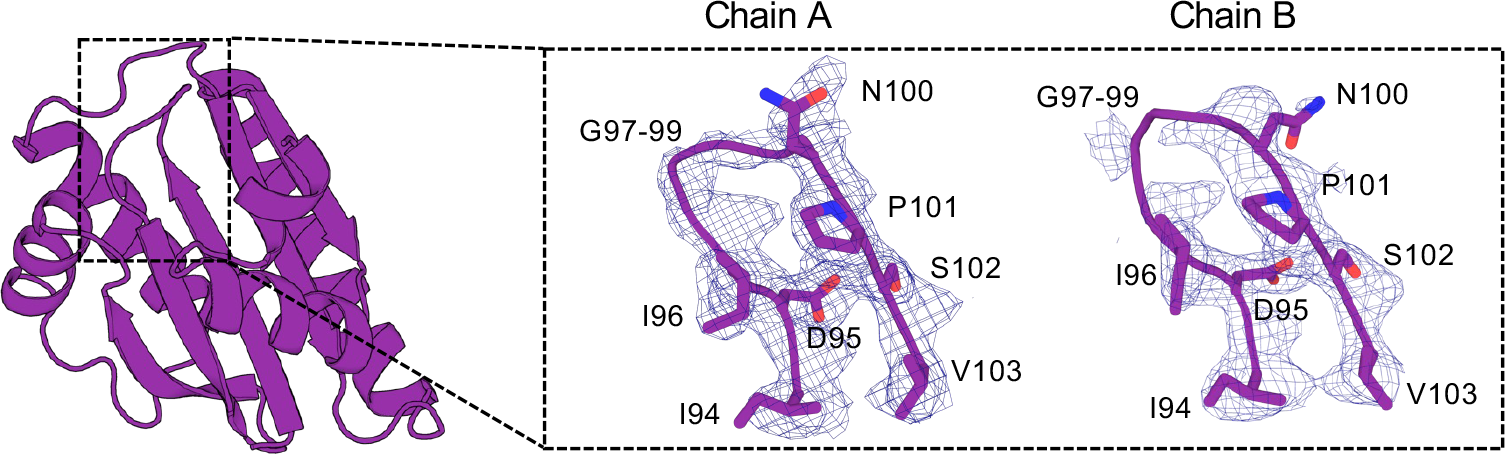
The glycine loop in the SEAL^GGG^ domain is highly flexible. Overview cartoon diagram of the SEAL structure with a detailed view of the “GGG loop” for both chains in the crystal asymmetric unit. Mesh represents the 2Fo-Fc map contoured at 1.0σ. The full loop on chain B is not resolved.

**Figure S9.**
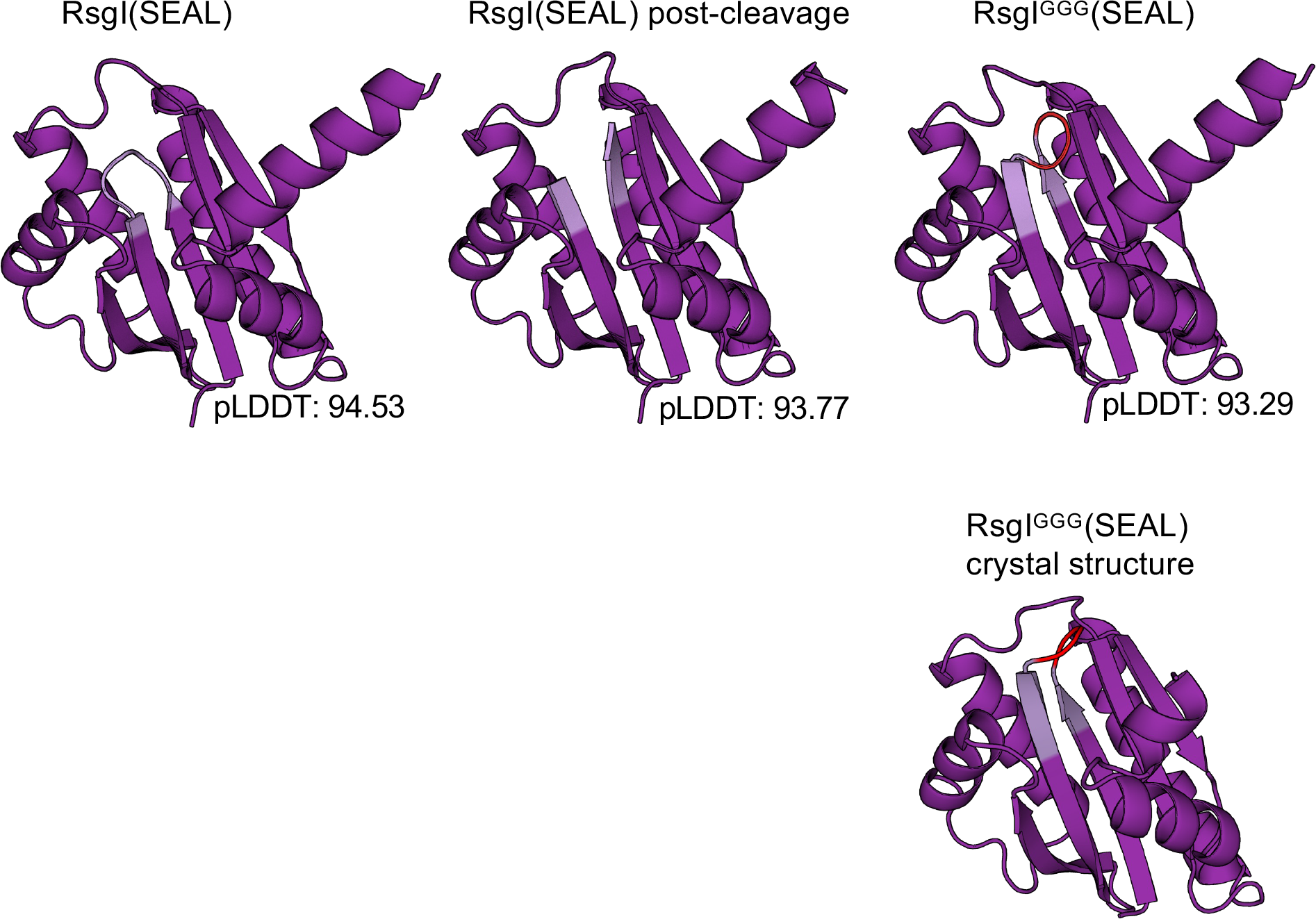
Alphafold2 predicts similar structures for autocleaved RsgI and RsgI^GGG^. AlphaFold2 predictions of uncleaved (left) and GGG variant (right) of RsgI’s SEAL domain compared to the AlphaFold-multimer prediction of the two cleaved fragments (middle). The conserved beta hairpin loop (DINPS) is colored in light purple and the inserted glycines colored in red. The cleaved SEAL domain and the GGG variant are both predicted to have extended ß- strands. This extension is confirmed in the SEAL^GGG^ crystal structure (bottom right).

**Figure S10.**
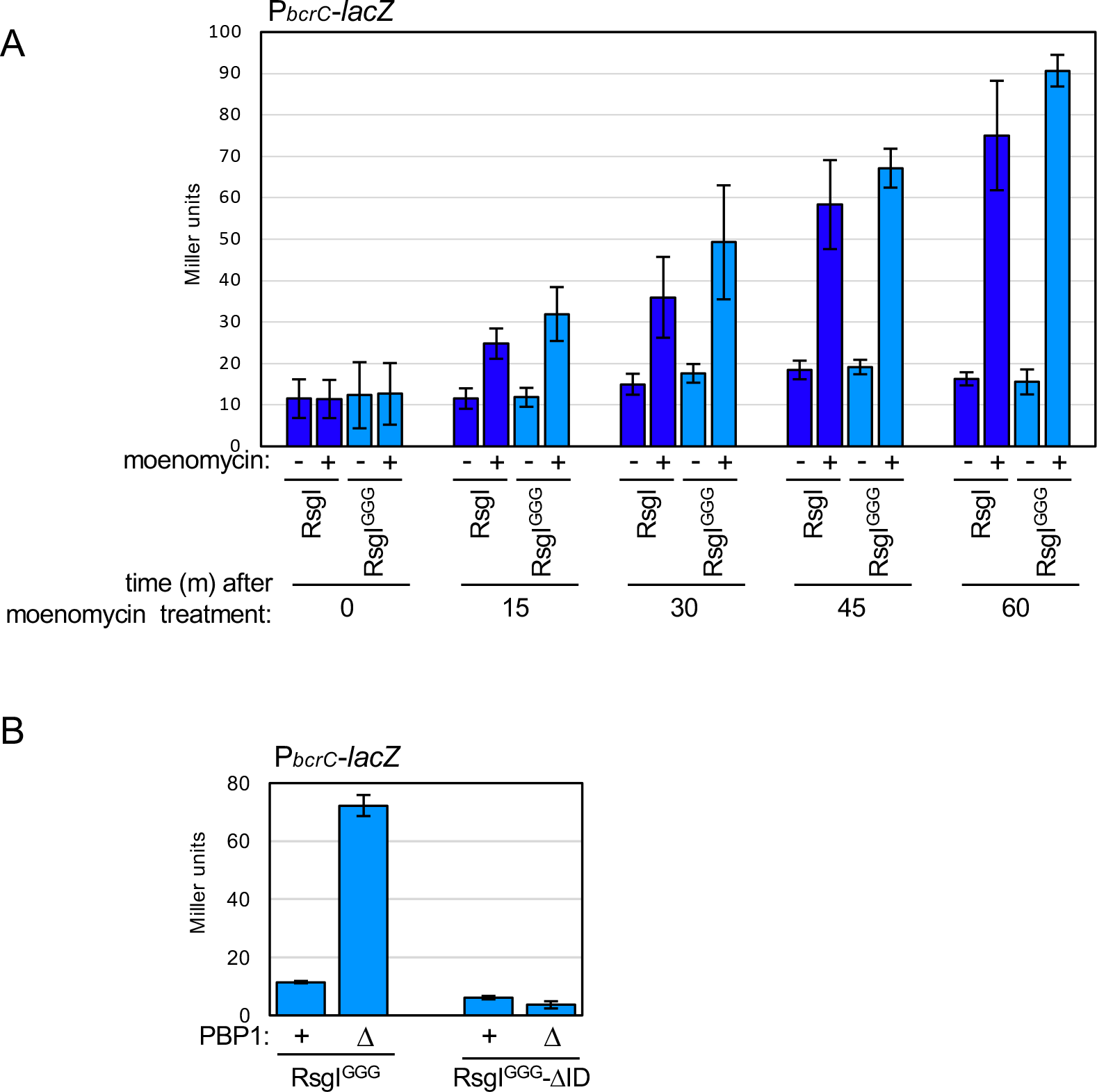
RsgI^GGG^ responds to cell wall defects in a manner that depends on its intrinsically disordered region. (A) Bar graph showing ß-galactosidase activity of a SigI-responsive (P*_bcrC_*) reporter before and after addition of moenomycin. Strains harboring untagged RsgI and RsgI^GGG^ respond similarly over a 60 min time course. **(B)** Bar graph showing beta-galactosidase activity of cells expressing RsgI^GGG^ or RsgI^GGG^ΔID in the presence and absence of PBP1. If RsgI^GGG^ lacks its intrinsically disordered region (ΔID), the cells are unable to respond to cell wall defects. All ß-galactosidase assays were performed in biological triplicate and error bars indicate standard error among them.

**Figure S11.**
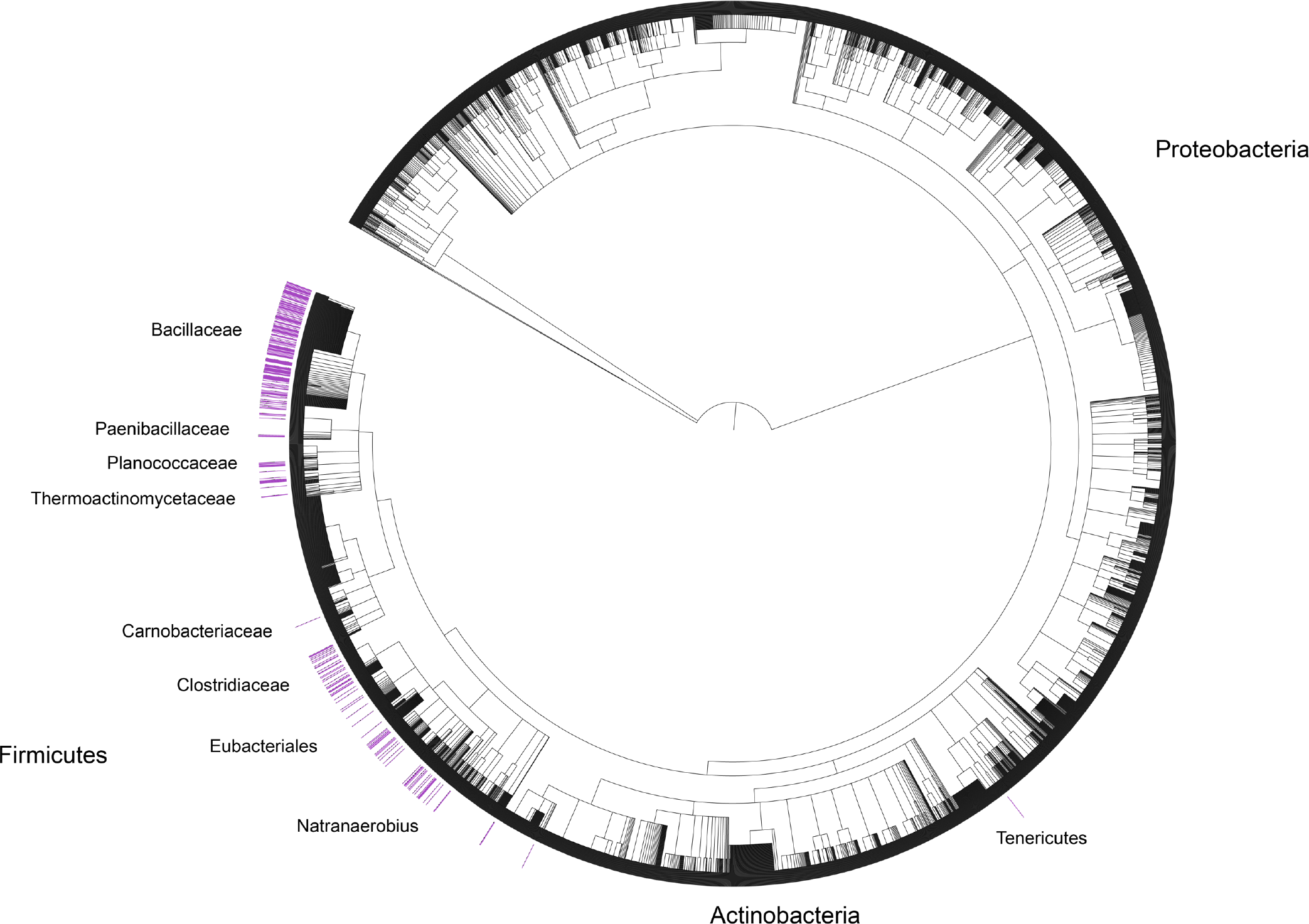
SEAL domains are broadly conserved among Firmicutes. Phylogenetic tree showing distribution of SEAL domains throughout 5767 bacterial taxa. The amino acid sequence of *B. subtilis* RsgI’s SEAL domain was used to query the Refseq database and the resulting species containing SEAL homologs are annotated in purple.

**Figure S12.**
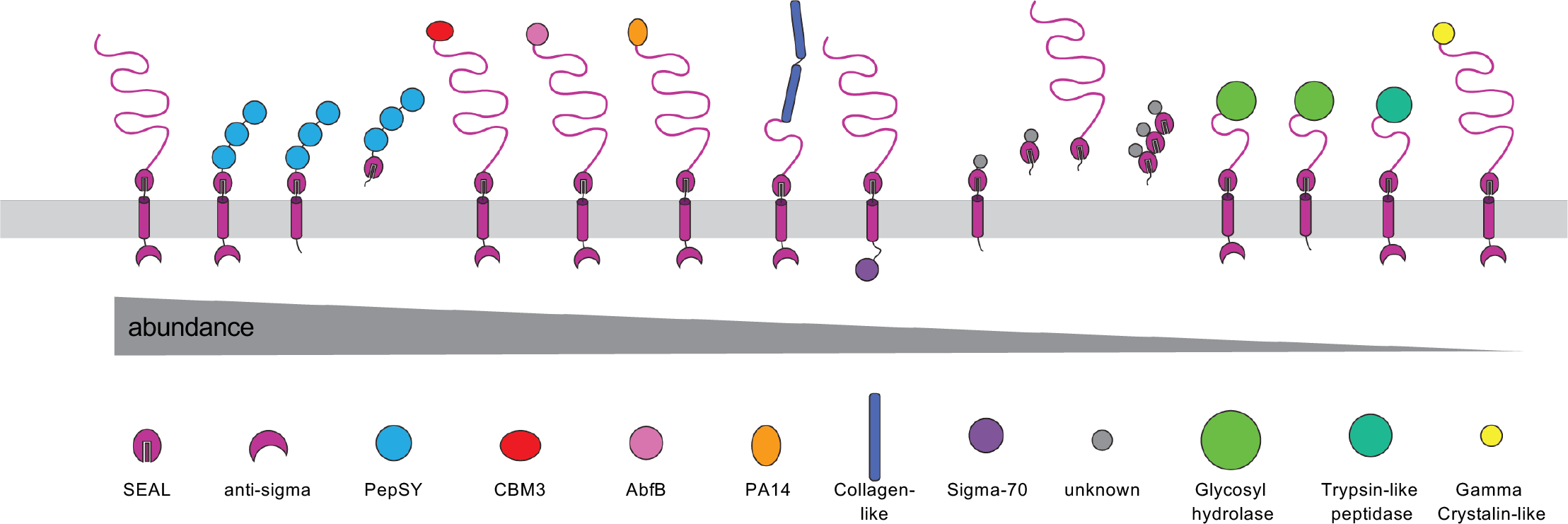
SEAL domains are found in proteins with diverse domain architectures. Schematics of proteins with SEAL domains found across Firmicutes. The amino acid sequence of *B. subtilis* RsgI’s SEAL domain was used to search the Refseq select protein database for sequence homologs. The resulting protein FASTA files were annotated for known Pfam domains using PfamScan. The protein architectures were reconstructed by binning the identified Pfam domains by protein accession number in the order identified. The protein families are displayed from most to least abundant. All SEAL-containing domains are predicted to be extracytoplasmic and either fused to a TM segment or a signal peptide.

**Figure S13.**
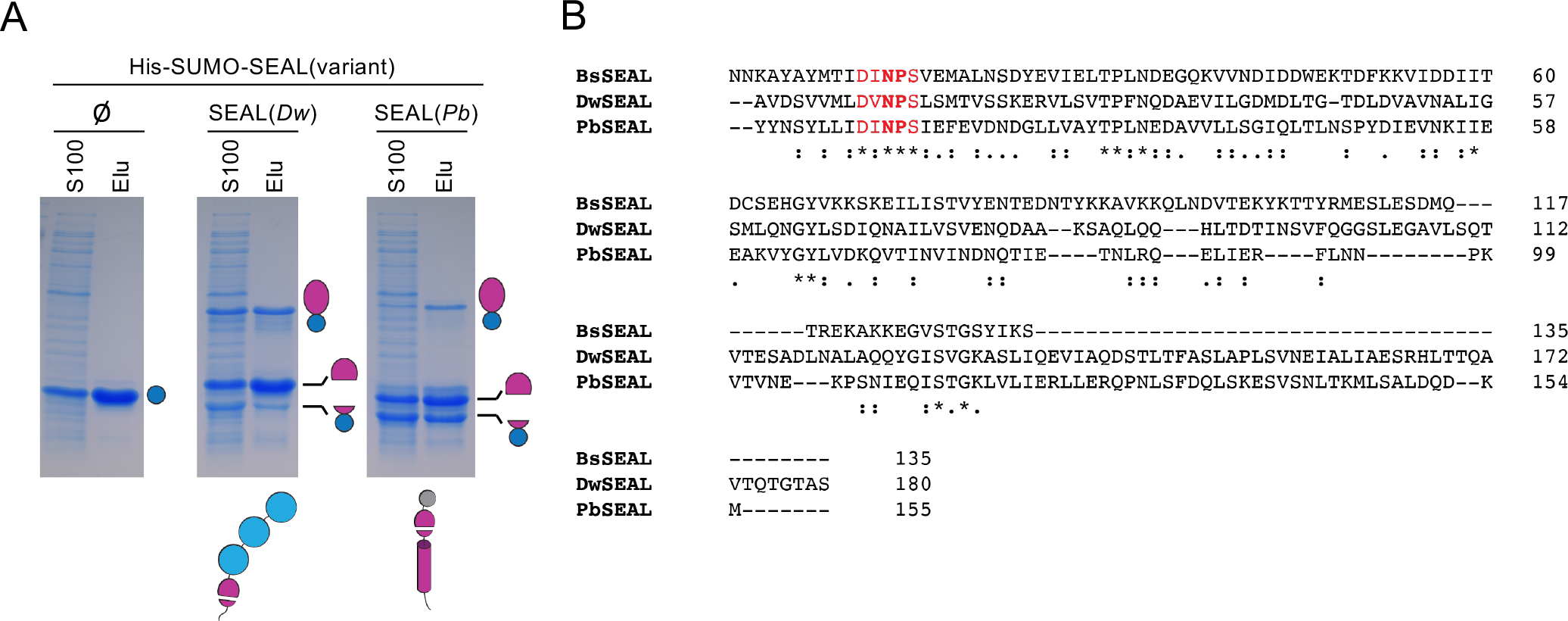
Autoproteolysis is a conserved feature of SEAL domains. (A) Coomassie-stained gels of His-SUMO(SEAL) homologs from *D. welbionis* (*Dw*) (EIO64_09220) and *P. brassicae* (*Pb*) (BN85316100) expressed and purified from *E. coli*. His-SUMO lacking a SEAL domain (ø) was included as a control. Schematics of full-length proteins are shown below the gels. The *B. welbionis* protein has a predicted signal peptide, a SEAL domain, and three putative PepSY domains. The *P. brassicae* protein has a TM segment, a SEAL domain, and a domain of unknown function. The domain is unannotated by Pfam but is conserved across homologs as a predicted six α-helical bundle with no characteristic sequence or structural homology. The *P. brassicae* protein lacks a sigma factor binding domain. **(B)** Multiple sequence alignment of *B. subtilis*, *D. welbionis*, and *P. brassicae* SEAL domains. The conserved beta-hairpin that is the site of autocleavage is highlighted in red, and the cleavage site is in bold.

## Supplementary Information

Supporting Text Plasmid Construction

**pAB47 [His-SUMO-RsgI-S99A (amp)]**

pAB47 was generated in a 2-piece isothermal assembly reaction using a PCR product amplified from pYB227 [amyE-sigI-rsgI (S99A) -S481-UTR (kan)(amp)] using oCH53 & oCH69 and pTD68 [His-SUMO (amp)] cut with BamHI.

pAB61 [yvbJ::PxylA-gfp-rsgI-His6 (erm)(amp)]

pAB61 was generated in a 2-piece isothermal assembly reaction using a PCR product amplified from PY79 gDNA with oAB112 & oAB113 and pYB200 [yvbJ::PxylA-gfp-His6 (erm)(amp)] cut with BamHI.

pAB66 [His-SUMO-RsgI(JMD)-S212A (amp)]

pAB66 was generated in a 2-piece isothermal assembly reaction using a PCR product amplified from pYB233[amyE-sigI-rsgI (S212A) -S481-UTR (kan)(amp)] using oCH53 & oCH69 and pTD68 [His-SUMO (amp)] cut with BamHI.

pAB67 [His-SUMO-RsgI-S99A-S212A (amp)]

pAB67 was generated in a 2-piece isothermal assembly reaction using a PCR product amplified from pYB227 [amyE-sigI-rsgI (S99A) -S481-UTR (kan)(amp)] using oCH53 & oAB117 and pTD68 [His-SUMO (amp)] cut with BamHI.

pAB78 [yvbJ::PxylA-gfp-rsgI-DI-GGG-NPS-His6 (erm)(amp)]

pAB78 was generated in a 3-piece isothermal assembly reaction using a PCR product amplified from PY79 gDNA with oAB112 & oAB128, a PCR product amplified using oAB113 & oAB129 and pYB200 [yvbJ::PxylA-gfp-His6 (erm)(amp)] cut with BamHI.

pAB79 [His-SUMO-RsgI(JMD)-DI-GGG-NPS (amp)]

pAB79 was generated in a 2-piece isothermal assembly reaction using a PCR product amplified from pAB78 with oCH53 & oCH69 and pTD68 [His-SUMO (amp)] cut with BamHI.

pAB87 [PT7-GFP-RsgI(JMD) (amp)]

pAB87 was generated in a 3-piece isothermal assembly reaction using a PCR product amplified from pAB61 using oAB141 & oAB144, a PCR product amplified from PY79 gDNA using oAB142 & oAB143 and pDHFR (NEB) cut with NdeI and BamHI.

pAB90 [amyE::sigI-rsgI-DI-GGG-NPS-S481UTR (kan)(amp)]

pAB90 was generated in a 3-piece isothermal assembly reaction using a PCR product amplified from PY79 gDNA using oAB35 & oAB128, a PCR product amplified from P79 gDNA using oAB36 & oAB129, and pYB225 cut with NheI.

pAB93 [His-SUMO-HtRsgI2(JMD) (amp)]

pAB93 was generated in a 2-piece isothermal assembly reaction using a PCR product amplified from RsgI2 GBlock using oAB154 & oAB155 and pTD68 [His-SUMO (amp)] cut with BamHI.

pAB101 [PT7-GFP-RsgI(JMD)-DI-GGG-NPS (amp)]

pAB101 was generated in a 3-piece isothermal assembly reaction using a PCR product amplified from pAB61 using oAB141 & oAB144, a PCR product amplified from pAB79 using oAB142 & oAB143, and pDHFR (NEB) cut with NdeI and BamHI.

pAB129 [amyE::sigI-rsgI-DI-GGG-NPS-ΔID-S481UTR (kan)(amp)]

pAB129 was generated in a 2-piece isothermal assembly reaction using a PCR product amplified from pAB90 using oAB35 & oYB425, and pYB225 cut with NheI.

pAB130 [yvbJ::PxylA-gfp-rsgI-DI-GGG-NPS-ΔID-His6 (erm)(amp)]

pAB130 was generated in a 2-piece isothermal assembly reaction using a PCR product amplified from pAB90 using oAB112 & oYB507, and pYB200 [yvbJ::PxylA-gfp-His6 (erm)(amp)] cut with BamHI.

pAB188 [His-SUMO-RsgI-GGG(JMD)-A88-S219 (amp)]

pAB188 was generated in a 2-piece isothermal assembly reaction using a PCR product amplified from pAB90 using oAB356 & oCH69, and pTD68 [His-SUMO (amp)] cut with BamHI.

**pAB214 [His-SUMO-SEAL(*Dwelbionis*) (amp)]**

pAB214 was generated in a 2-piece isothermal assembly reaction using a PCR product amplified from SEAL(D.welbionis) GBlock using oAB410 & oAB411, and pTD68 [His-SUMO (amp)] cut with BamHI.

**pAB216 [His-SUMO-SEAL(*Pbrassicae*) (amp)]**

pAB216 was generated in a 2-piece isothermal assembly reaction using a PCR product amplified from SEAL(Pbrassicae) using oAB412 & oAB413, and pTD68 [His-SUMO (amp)] cut with BamHI.

pCH26 [His-SUMO-CtRsgI(JMD) (amp)]

pCH26 was generated in a 2-piece isothermal assembly reaction using a PCR product amplified from *C. thermocaliphilum* gDNA using oCH63 & oCH54, and pTD68 [His-SUMO (amp)] cut with BamHI.

pCH27 [His-SUMO-HtRsgI4(JMD) (amp)]

pCH27 was generated in a 2-piece isothermal assembly reaction using a PCR product amplified from *H. thermocellum* gDNA using oCH65 & oCH66, and pTD68 [His-SUMO (amp)] cut with BamHI.

pCH29 [His-SUMO-BsRsgI(JMD) (amp)]

pCH29 was generated in a 2-piece isothermal assembly reaction using a PCR product amplified from PY79 gDNA using oCH53 & oCH69, and pTD68 [His-SUMO (amp)] cut with BamHI.

### Supplemental Tables

**Table S1.** Hits from the DALI server search with Alphafold predicted *Bacillus subtilis* SEAL

|  | *Description* | *Organism* | *Chain* | *Z score* | *RMSD* | *Length* | *#Resi* | *% ID* |
| --- | --- | --- | --- | --- | --- | --- | --- | --- |
| 1 | Protein N-terminal asparagine amidohydrolase | *Homo sapiens* | 6a0e-A | 6.1 | 2.7 | 85 | 305 | 5 |
| 2 | YlmD (cysteine hydrolase) | *Geobacillus stearothermophilus* | Lt8h-A | 5.6 | 2.7 | 74 | 272 | 9 |
| 3 | APOBEC3H (mRNA editing enzyme) | *Macaca nemestrina* | 5w3v-A | 5.6 | 3.2 | 83 | 186 | 5 |
| 4 | Deep network hallucinated protein | De novo synthetic construct | 7m0q-B | 5.5 | 2.8 | 79 | 97 | 6 |
| 5 | APOBEC3H (mRNA editing enzyme) | *Macaca mulatta* | 6p40-A | 5.5 | 6 | 91 | 364 | 3 |
| 6 | TOP7 surface mutant | De novo synthetic | 7fao-A | 5.5 | 3 | 76 | 96 | 13 |
| 7 | De novo designed protein | De novo synthetic | 2mra-A | 5.4 | 3 | 84 | 117 | 15 |
| 8 | NTPDase 2 (nucleoside triphosphate hydrolase) | *Rattus norvegicus* | 4br5-A | 5.4 | 3.7 | 110 | 422 | 4 |
| 9 | De novo designed beta sheet | De novo synthetic | 4kyz-A | 5.3 | 5 | 82 | 167 | 11 |
| 10 | VASH2/SVBP (tubulin-tyrosine carboxypeptidase) | *Homo sapiens* | 6jze-A | 5.3 | 3.9 | 76 | 254 | 7 |

**Table S2.** Summary of data collection, phasing and refinement statistics

|  | RsgI<br>(8T9N) |
| --- | --- |
| **Data collection** |  |
| Space group | P 1 2<sub>1</sub> 1 |
| Cell dimensions |  |
| *a*, *b*, *c* (Å) | 38.175, 63.889, 59.935 |
| $\alpha$, $\beta$, $\gamma$ (°) | 90.0, 92.6, 90.0 |
| Resolution (Å) | 38.13–1.90 (1.94–1.90) |
| *R*<sub>pim</sub> | 3.8 (87.9) |
| *I* / $\sigma(I)$ | 10.1 (1.0) |
| Completeness (%) | 98.6 (97.9) |
| Redundancy | 3.7 (3.5) |
| **Refinement** |  |
| Resolution (Å) | 38.13–1.90 |
| No. reflections |  |
| Total | 82070 |
| Unique | 22424 |
| Free | 1994 |
| *R*<sub>work</sub> / *R*<sub>free</sub> | 23.03 / 27.48 |
| No. atoms |  |
| Protein | 1983 |
| Ligand / ion | – |
| Water | 80 |
| *B*-factors |  |
| Protein | 62.65 |
| Ligand / ion | – |
| Water | 54.73 |
| R.m.s. deviations |  |
| Bond lengths (Å) | 0.004 |
| Bond angles (°) | 0.688 |
| Ramachandran<br>(favoured/allowed) | 97.92 / 0.88 |
| Rotamer Outliers | 0.88 |
| MolProbity Score | 1.60 |

\*Dataset was collected from individual crystals.
\*\*Values in parentheses are for the highest resolution

**Table S3.** SEAL PfamScan results

| **Pfam** | **Name** | **Count** |
| --- | --- | --- |
| PF12791 | Anti-sigma factor N-terminus | 2914 |
| PF03413 | Peptidase propeptide and YpeB (PepSY) domain | 229 |
| PF00942 | Cellulose binding domain (CBM3) | 11 |
| PF05270 | Alpha-L-arabinofuranosidase B (AbfB) domain | 10 |
| PF07691 | PA14 Domain | 7 |
| PF01391 | Collagen triple helix repeat | 6 |
| PF13365 | Trypsin-like peptidase domain | 5 |
| PF04542 | Sigma-70 region 2 | 3 |
| PF00030 | beta/gamma crystallin-like | 1 |
| PF00331 | Glycosyl hydrolase family 10 | 1 |

**Table S4.** Strains used in this study

| **Strain** | **Background** | **Genotype** | **Reference** | **Figure** |
| --- | --- | --- | --- | --- |
| BAB325 | *B.subtilis*<br>PY79 | *yvbJ::PxylA-gfp-RsgI-His (erm) ΔrsgI::spc* | This Study | 5 |
| BAB331 | *B.subtilis*<br>PY79 | *yvbJ::PxylA-gfp-RsgI-DI-GGG-NPS-His (erm) ΔrsgI::spc* | This Study | 5, 6 |
| BAB332 | *B.subtilis*<br>PY79 | *yvbJ::PxylA-gfp-RsgI-His (erm) ΔrsgI::spc ΔponA::kan* | This Study | 5 |
| BAB338 | *B.subtilis*<br>PY79 | *yvbJ::PxylA-gfp-RsgI-DI-GGG-NPS-His (erm) ΔrsgI::spc ΔponA::kan* | This Study | 5, 6 |
| BAB19 | *B.subtilis*<br>PY79 | *sacA::Pveg-gfp ( Phleo) yhdG::PbcrC-optRBS-lacZ (cat) Δ(sigI-rsgI)::spc amyE::sigI-rsgI (kan)* | This Study | 5, S10 |
| BAB29 | *B.subtilis*<br>PY79 | *sacA::Pveg-gfp ( Phleo) yhdG::PbcrC-optRBS-lacZ (cat) Δ(sigI-rsgI)::spc amyE::sigI-rsgI (kan) ΔponA::erm* | This Study | 5, S10 |
| BAB321 | *B.subtilis*<br>PY79 | *sacA::Pveg-gfp ( Phleo) yhdG::PbcrC-optRBS-lacZ (cat) Δ(sigI-rsgI)::spc amyE::sigI-rsgI-DI-GGG-NPS (kan)* | This Study | 5, S10 |
| BAB323 | *B.subtilis*<br>PY79 | *sacA::Pveg-gfp ( Phleo) yhdG::PbcrC-optRBS-lacZ (cat) Δ(sigI-rsgI)::spc amyE::sigI-rsgI-DI-GGG-NPS (kan) ΔponA::erm* | This Study | 5, S10 |
| BAB443 | *B.subtilis*<br>PY79 | *sacA::Pveg-gfp ( Phleo) yhdG::PbcrC-optRBS-lacZ (cat) Δ(sigI-rsgI)::spc amyE::sigI-rsgI-DI-GGG-NPS-ΔID (kan)* | This Study | S10 |
| BAB447 | *B.subtilis*<br>PY79 | *sacA::Pveg-gfp ( Phleo) yhdG::PbcrC-optRBS-lacZ (cat) Δ(sigI-rsgI)::spc amyE::sigI-rsgI-DI-GGG-NPS-ΔID (kan) ΔponA::erm* | This Study | S10 |
| BAB450 | *B.subtilis*<br>PY79 | *yvbJ::PxylA-gfp-RsgI-DI-GGG-NPS-ΔID-His (erm) ΔrsgI::spc* | This Study | 6 |
| BAB453 | *B.subtilis*<br>PY79 | *yvbJ::PxylA-gfp-RsgI-DI-GGG-NPS-ΔID-His (erm) ΔrsgI::spc ΔponA::kan* | This Study | 6 |
| BAB339 | *B.subtilis*<br>PY79 | *yvbJ::PxylA-gfp-RsgI-His (erm) ΔrsgI::spc ΔrasP::tet* | This Study | 6 |
| BAB345 | *B.subtilis*<br>PY79 | *yvbJ::PxylA-gfp-RsgI-DI-GGG-NPS-His (erm) ΔrsgI::spc ΔrasP::tet* | This Study | 6 |

**Table S5.**
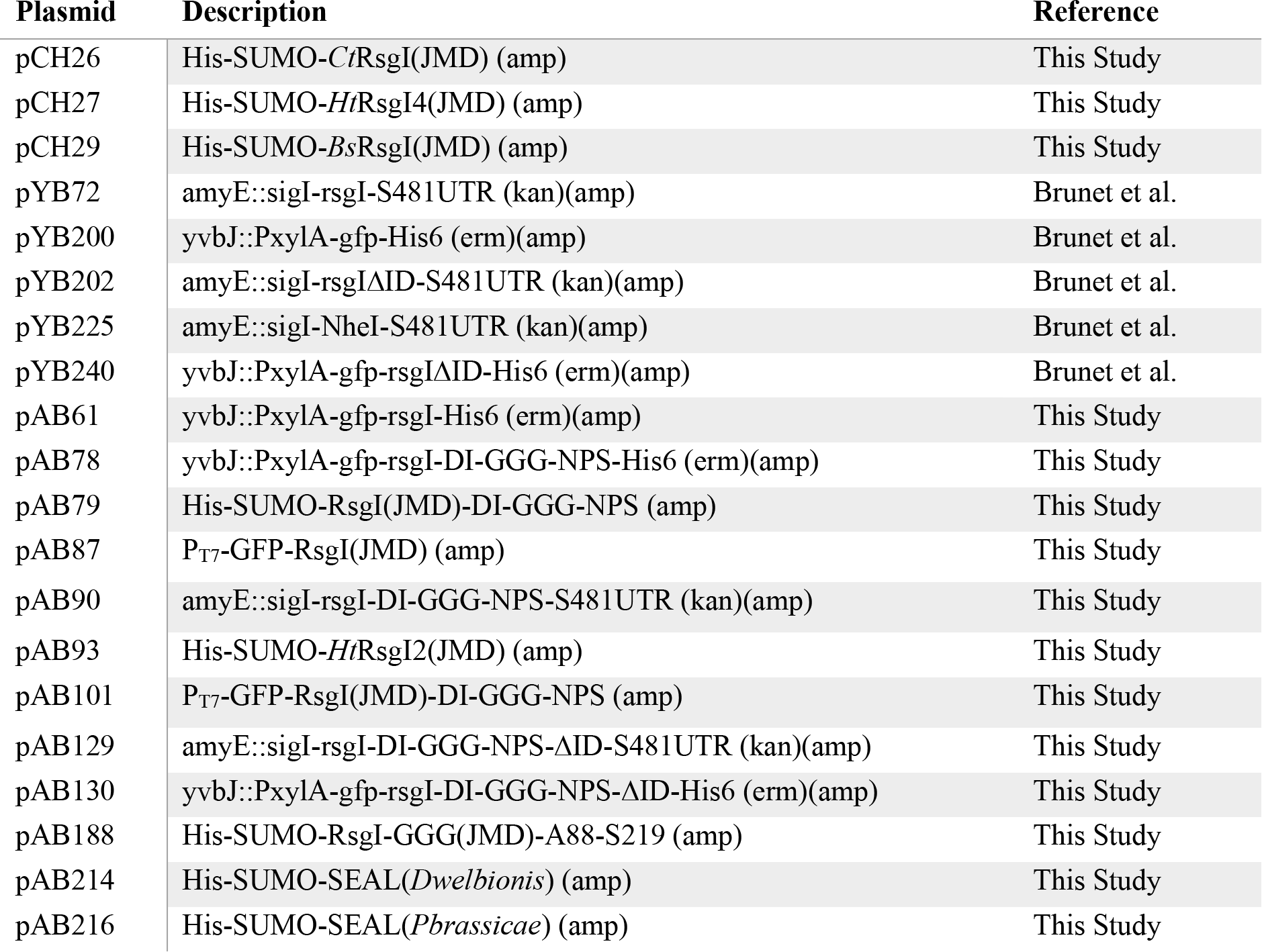
Plasmids used in this study

| **Plasmid** | **Description** | **Reference** |
| --- | --- | --- |
| pCH26 | His-SUMO-*CtRsgI*(JMD) (amp) | This Study |
| pCH27 | His-SUMO-*HtRsgI4*(JMD) (amp) | This Study |
| pCH29 | His-SUMO-*BsRsgI*(JMD) (amp) | This Study |
| pYB72 | amyE::sigI-rsgI-S481UTR (kan)(amp) | Brunet et al. |
| pYB200 | yvbJ::PxylA-gfp-His6 (erm)(amp) | Brunet et al. |
| pYB202 | amyE::sigI-rsgIΔID-S481UTR (kan)(amp) | Brunet et al. |
| pYB225 | amyE::sigI-NheI-S481UTR (kan)(amp) | Brunet et al. |
| pYB240 | yvbJ::PxylA-gfp-rsgIΔID-His6 (erm)(amp) | Brunet et al. |
| pAB61 | yvbJ::PxylA-gfp-rsgI-His6 (erm)(amp) | This Study |
| pAB78 | yvbJ::PxylA-gfp-rsgI-DI-GGG-NPS-His6 (erm)(amp) | This Study |
| pAB79 | His-SUMO-RsgI(JMD)-DI-GGG-NPS (amp) | This Study |
| pAB87 | P<sub>T7</sub>-GFP-RsgI(JMD) (amp) | This Study |
| pAB90 | amyE::sigI-rsgI-DI-GGG-NPS-S481UTR (kan)(amp) | This Study |
| pAB93 | His-SUMO-*HtRsgI2*(JMD) (amp) | This Study |
| pAB101 | P<sub>T7</sub>-GFP-RsgI(JMD)-DI-GGG-NPS (amp) | This Study |
| pAB129 | amyE::sigI-rsgI-DI-GGG-NPS-ΔID-S481UTR (kan)(amp) | This Study |
| pAB130 | yvbJ::PxylA-gfp-rsgI-DI-GGG-NPS-ΔID-His6 (erm)(amp) | This Study |
| pAB188 | His-SUMO-RsgI-GGG(JMD)-A88-S219 (amp) | This Study |
| pAB214 | His-SUMO-SEAL(*Dwelbionis*) (amp) | This Study |
| pAB216 | His-SUMO-SEAL(*Pbrassicae*) (amp) | This Study |

**Table S6.**
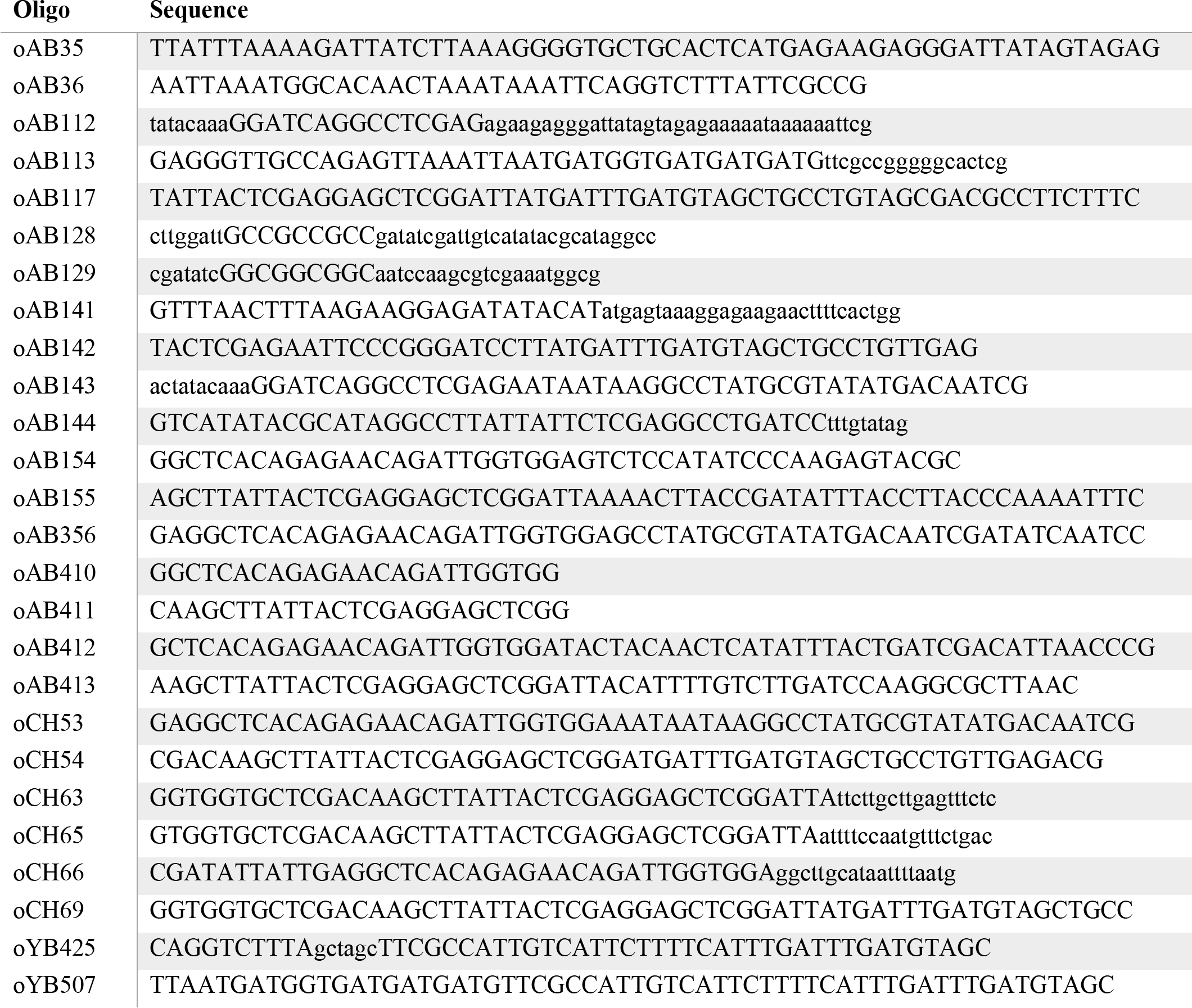
Oligonucleotides used in this study

**Table S7.**
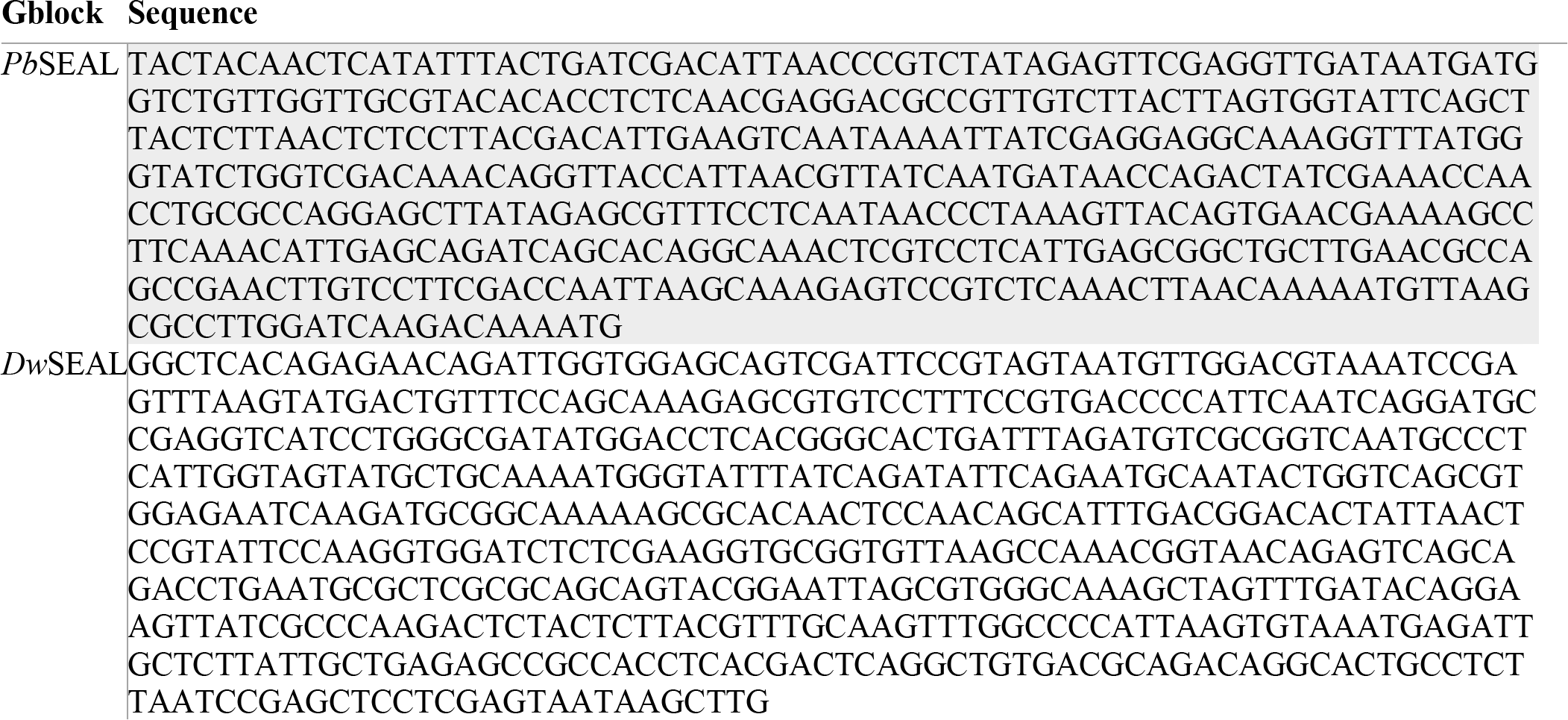
Gene blocks used in this study

## Notes

### Competing Interest Statement

The authors have declared no competing interest.

